# Viral Phrenology

**DOI:** 10.1101/2021.09.02.458596

**Authors:** David Wilson, Danielle Roof

## Abstract

We introduce Viral Phrenology, a new scheme for understanding the genomic composition of spherical viruses base on the locations of their structural protrusions. We used icosahedral point arrays to classify 135 distinct viral capsids collected from over 600 capsids available in the VIPERdb. Using gauge points of point arrays, we found 149 unique structural protrusions. We then show how to use the locations of these protrusions to determine the genetic composition of the virus. We then show that ssDNA, dsDNA, dsRNA and ssRNA viruses use different arrangements for distributing their protrusions. We also found that Triangulation number is also partially dependent on the structural protrusions. This analysis begins to tie together Baltimore classification and Triangulation number using point arrays.

## 1. Introduction

We studied 135 distinct spherical viruses taken from VIPERdb [1] and found that the locations of their structural protrusions often indicated their genetic composition. We refer to this method of determining a virus’s genetic composition by examining the placement of its protrusions as Viral Phrenology. The structural protrusions were found by classifying the viruses using our modified fitting methods [2] for icosahedral point arrays [3,4]. Point arrays provide highly specific geometric constraints on the arrangement of viral proteins, and all spherical viruses studied so far conform to one or more of these arrays [2–13]. We have previously shown that all protruding features of spherical viruses are located on the gauge points of these arrays, though it was not yet known that these locations also indicated the genomic composition. The gauge points determine the overall radial scaling of a point array and are all located on the 15 icosahedral great circles which subtend neighboring symmetry axes in the asymmetric unit [2,14]. We also show that not all gauge points are used for both DNA and RNA viruses. Additionally we see that the Triangulation number [15] is also semi-indicated by the location of protrusions.

Spherical virus capsids have icosahedral symmetry, and are classified by the Triangulation (T)-number, which posits that viral capsid proteins are arranged in nearly identical chemical environments, known as quasi-equivalence. In general, T-number specifies the number of identical capsid proteins (60T), that there are 12 pentameric units and 10(*T* −1) hexameric units. There are a number of viruses that make exceptions to these rules, though remarkable they still ascribe to the architectures prescribed by the T-number. For example, SV40 [16] has the architecture of T7d, though it is only composed of 360 proteins, rather than 420 and it is made entirely from pentamers. Remarkably, the virus maintains icosahedral symmetry with pentamers located at each hexamer location. In addition, there are pseudo-T (pT) number viruses, which mimic larger T-number architectures using either different proteins or fewer multi-domain proteins [17]. For example, Human Rhinovirus 16 is a pT3 virus that is composed of only 60 proteins, yet it has a T3 architecture.

While spherical viruses have been well described by T-number, this classification does not provide any connection to the Baltimore Classification system [18], which organizes viruses into seven groups based on how they use mRNA in their replication cycle [19]. While cellular life is restricted to storing its genetic information within dsDNA, viruses are much more diverse. Viruses utilize dsDNA (BC I & BC VII), ssDNA (BC II), dsRNA (BC III) and ssRNA (BC IV, V & VI) genomes. To our knowledge, this study is the first to reveal the connection between the geometric arrangement of capsid proteins and the genetic composition of the virus. Given that the restrictions of point arrays are so specific, this understanding could inform evolution and mutation studies, development of virus-like particles (VLPs), and other potential therapeutic virus treatments.

## 2. Materials and Methods

We constructed a library of 135 distinct spherical capsids and then determined their best fitting point arrays. We then combined the gauge points, which correspond to the key structural protrusions, with the T-number and genomic composition of the viruses.

### 2.1. Virus Library

We gathered over 600 spherical virus structures from those deposited in VIPERdb [1]. We then narrowed this list down, by keeping each unique family, genus, Triangulation number and Baltimore Classification combination. We then chose from the remaining structures the ones with the best resolution; when ties existed, we kept both structures, see Appendix A for the complete library. The result of this filtering lead to 135 spherical virus capsids, ranging from T1 to T169 and also included pseudo T-numbers. The genomic composition and T-number for this library is shown in Table 1. This criteria also included some viruses structures which were bound small molecules or complexed with antibodies. The goal of this methodology was to create a diverse data set, and in total there are 50 distinct Virus families and 105 distinct genera. We decided to focus only on the genomic composition, rather than the mRNA pathways specified for simplicity. To our knowledge, there are no spherical viruses with BC V or BC VI in the VIPERdb [1], so in general when we say dsDNA we mean BC I, ssDNA is BC II, etc. The only ambiguity exists for circular-dsDNA genome with an RNA-phase, as in HBV which is BC VII. Lastly, not all capsids found in VIPERdb [1] had an identified genome.

**Table 1.** Viral Capsid Data set. We analyzed 135 distinct icosahedral capsids found in the VIPERdb [1]. Here we present their genomic composition vs T-number to illustrate the diversity of capsids and their genomes. There are 10 capsids not shown here, pT21(2), pT25(2), pT27, T28d, pT31, T43, pT169, T169, which were all dsDNA genomes. We note that nearly all of ssDNA capsids are T1 (87%), and most of ssRNA capsids (70%) are T3 or pT3. Most of T3 capsids are composed only of ssRNA (87%). T7, T16 and larger capsids were found to only contain dsDNA. In summary, knowing the T-number only provides limited information on the genome, as does only knowing the genome provide little information on the T-number. We did not find any T12 or T19 capsids as might be expect based on [20], though pT27 and T169 were present.

| Genome | T1 | T2 | T3 | pT3 | T4 | T7d | T7l | T9 | T13 | T16 | Total |
| --- | --- | --- | --- | --- | --- | --- | --- | --- | --- | --- | --- |
| dsDNA | 2 | 1 | 2 | 0 | 2 | 5 | 11 | 0 | 2 | 3 | 28 |
| ssDNA | 13 | 0 | 1 | 0 | 0 | 0 | 0 | 0 | 1 | 0 | 15 |
| dsRNA | 7 | 4 | 0 | 1 | 0 | 0 | 0 | 0 | 5 | 0 | 17 |
| ssRNA | 13 | 0 | 27 | 16 | 4 | 0 | 0 | 1 | 0 | 0 | 61 |
| None | 1 | 0 | 1 | 0 | 1 | 0 | 0 | 1 | 0 | 0 | 4 |
| Totals | 36 | 5 | 31 | 17 | 7 | 5 | 11 | 2 | 8 | 3 | 125* |

### 2.2. Point Arrays

We classify viruses according to their best fitting icosahedral point array(s) [3,4] using our fitness criteria introduced in [2,14]. These point arrays prescribe specific geometric constraints on viral capsids and their genetic material. These constraints are radially-distributed representations of icosahedral rotation symmetry which set limits on the angular orientation and distribution of virus capsid proteins, see Figure 1. While in principle, viruses could conform to multiple point arrays, in practice we find most viruses conform to only one or two arrays [2]. This classification is powerful, as it indicates geometric locations where viruses need to be modified with care, as well as other potential geometric locations where modifications should be relatively easy to accomplish. This classification is purely geometric in nature and does not take into account local surface chemistry, steric hindrance, or solvent accessibility.

**Figure 1.**
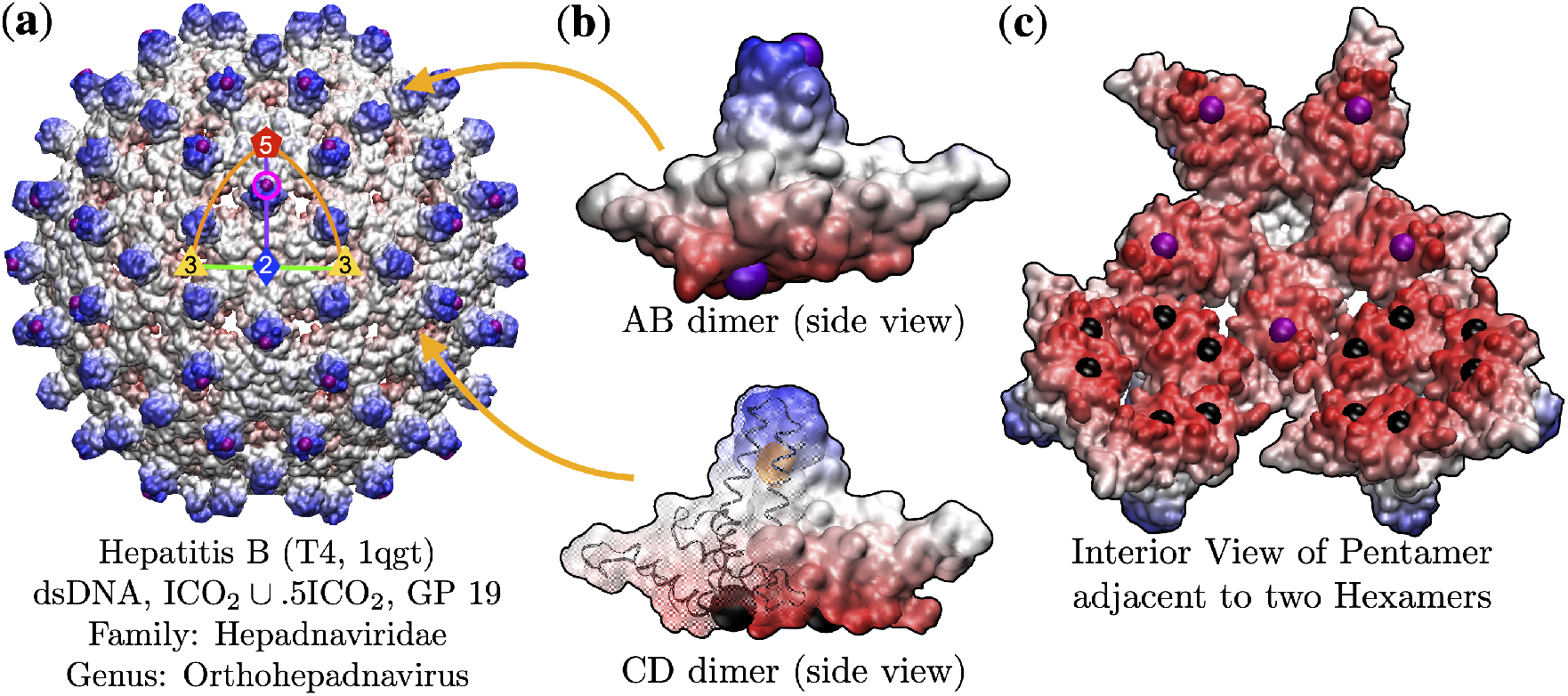
(**a**) The Hepatitis B capsid (HBV, T4, 1qgt [21]) is composed of 240 chemically-identical proteins (120 dimers) arranged with icosahedral symmetry. This arrangement leads to 12 pentamer and 30 hexamer subunits. There are no covalent bonds between any of these proteins, therefore weak van der Waals interactions are dominant. The AB dimers protrude between the 5 and 2 fold axes at gauge point 19 (pink circle). This gauge point determines nearly all of the other interior points, indicating that having protruding features at these locations puts constraints on the entire interior structure. None of these constraints are required by icosahedral symmetry. Gauge point 19 imposes inflexible restrictions on the choice of point arrays, either ICO_2_ or IDD_5_, are the only two point arrays available to this gauge point [2,14]. These are known as sister arrays, and are identical except for a single radial level and without more coordinate information on the genome, are indistinguishable at this level [2]. Each of these point arrays specify restrictions on three of the four protein chains A, C & D. (**b**) The capsid proteins pair up to form two dimers (AB & CD) which molecular dynamics simulations have shown to have different vibrational frequencies [22]. This is consistent with the point arrays, as the AB dimer is constrained at the top and bottom, and the CD dimer is constrained at the middle and bottom. We have highlighted the C-chain protein backbone, which is constrained at two points to each side of the point array element (orange). (**c**) The interior view of a pentamer and two hexamer subunits. We see that the point arrays put constraints on the interior of the hexamer (black) and pentamer (purple). The two point arrays available to GP 19, ICO_2_ and IDD_5_ specify all the points shown in **(a)** – **(c)**, except the purple points seen the interior pentamer in **(c)**. These purple points comes from the union of ICO_2_ ~ .5ICO_2_. This array imposes a restriction on chain B and is therefore the best fitting point array. There is not a similar union available for IDD_5_. It is important to note that none of the points of point array ICO_2_ ~ .5ICO_2_ can be removed or moved, there is only 1 degree of freedom, the overall radial scaling [2].

Icosahedral point arrays offer a novel tool for analyzing viruses [3,4,14]. We have shown previously that amino acids near these geometric locations are likely critical to capsid stability and that many modifications, from amino acid substitution to ligand attachment are either prohibited or requires more care than expected by standard analysis such as steric hindrance and solvent accessibility [2]. Point arrays are constructed by applying an affine extension to a base icosahedron (ICO), dodecahedron (DOD) or icosadodecahedron (IDD) along the 2-, 3- or 5-fold symmetry axes direction then re-applying icosahedral symmetry. Each affine extension is characterized by the scale length of the base, in terms of the golden ratio *ϕ* and there are 55 different single base arrays [2–4]. For example, 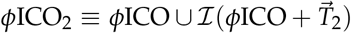, where ℐ represents applying the 60 rotations of the icosahedral group; This point array has a single gauge point, GP 21. Each array has a single free-parameter, the overall radial scaling. Point arrays with the same affine extension can be combined to form larger point arrays [2–4]. Many of our virus fits combine multiple point arrays.

We have shown previously that all spherical protrusions are found on, or very close to the gauge points [14]. Our point array classification scheme begins by determining the center of mass for each viral protrusion, and then determining the gauge points nearest to them, based on an angular cutoff [14], see Figure 2. We then scale all point arrays associated with these gauge points and optimize their RMSD fit [2]. In principle, a virus may have multiple protruding features on different gauge points. We refer to protruding features with gauge points that correspond to best fit point arrays as structural protrusions; that is their specific locations determine almost all of the internal structural constraints. The location of the gauge points determine the radial size of the point array and limit the possible interior arrays, imposing strong restrictions on the relative placement of capsid proteins at each radial level [2], see Figure 2. It is also important to note, each gauge point not located on an icosahedral symmetry axes (2,3,5) has only 2 point arrays associated with it, so the locations of the gauge points place strong restrictions on the arrangement of capsid proteins. Our RMSD measure is found by

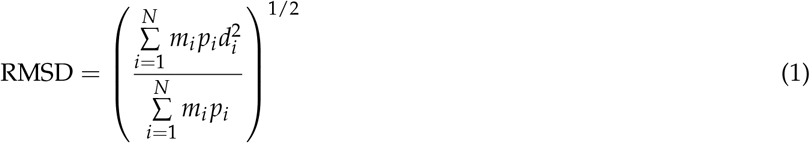

where *d*_*i*_ is the minimum distance from the *i*^th^ point array element to the nearest protein surface(s) or genome (if available), *p*_*i*_ is the number of different proteins near the point *i* (*e.g*. 5 for a point on a 5-fold axes or 2 if two proteins are equidistant from the same point). Finally *m*_*i*_ is the number of times the point appears in the full point array (*i.e*., 12, 20, 30 or 60) and *N* is the total number of elements in the point array.

**Figure 2.**
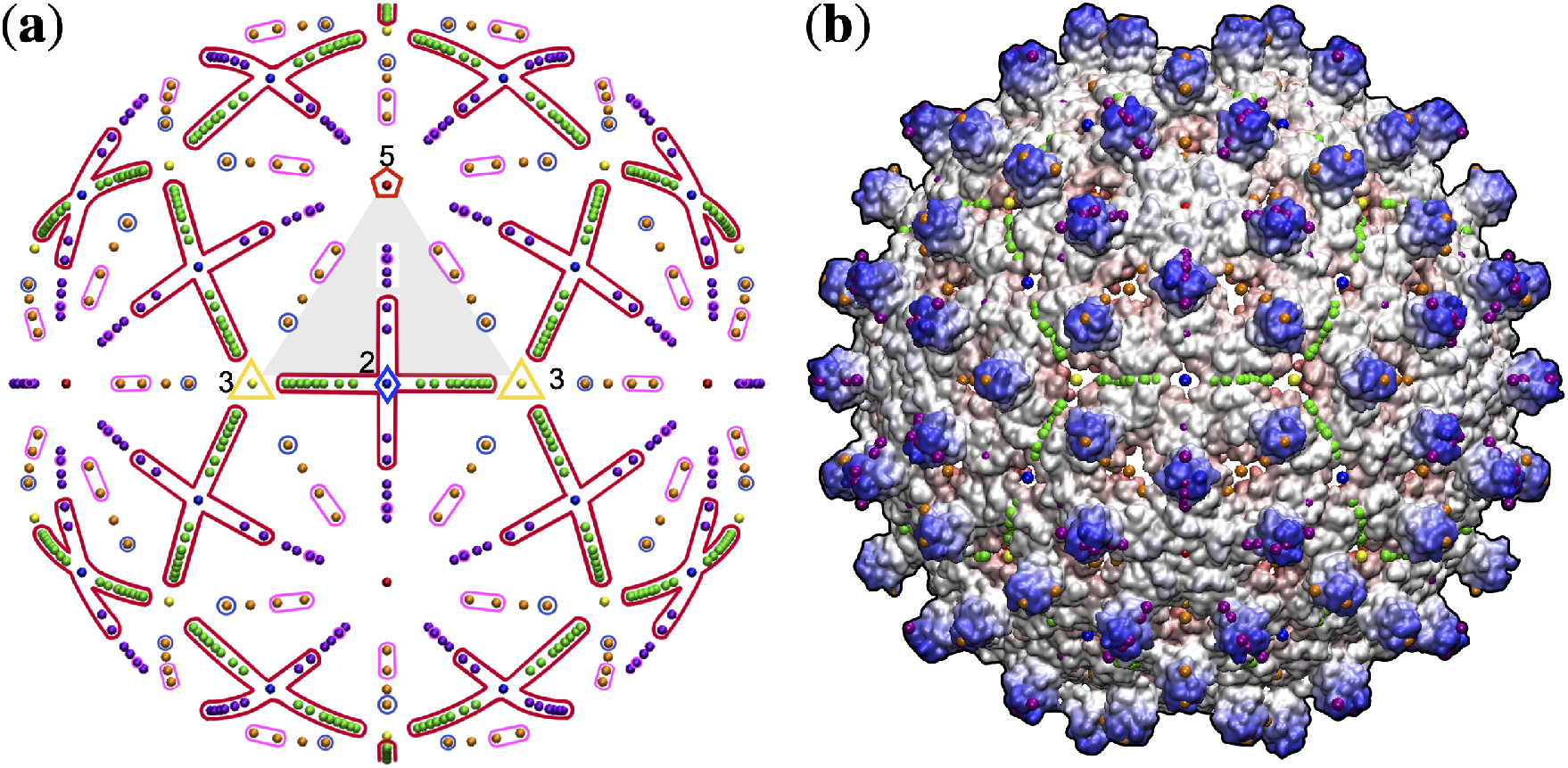
(**a**) The gauge points of the point arrays [2–4,14] are found on the icosahedral great circles which subtend neighboring symmetry axes, and define the Asymmetric Unit (AU), shown in grey. The AU is a 1/60^*th*^ representative section of the full icosahedral capsid and there are 21 unique gauge points within it. These points are all colored based on where they are relative to the symmetry axes and the great circles they connect; 5-folds are red, 3-folds are yellow, 2-folds are blue, then purple is between 5 (red) and 2 (blue), green is between 2 (blue) and 3 (yellow), orange is between 5 (red) and 3 (yellow), finally black points are not on great circles, instead they are found anywhere else and are referred to as bulk points. The angular locations of these points are shown to scale. By analyzing the structural protrusions and the genomic composition of viruses, we found that only RNA viruses are found to use the encircled red region (⊥), and also dominate areas encircled with pink. Finally, only ssDNA viruses use the encircled blue region (near 3-fold axes). (**b**) The gauge points arranged atop HBV (T4, 1qgt) virus. There are two potential structural protrusions within the AU, located along the 5-2 arc (purple) and the 3-5 arc (orange). We find that only the point arrays corresponding to purple protrusion GP 19 have corresponding arrays which meet the criteria for a proper fit [2], so there is only one structural protrusion in HBV.

While these protrusions are known to play several important biological functions in gaining entry to new cells, the fact that they also constrain the structure of the interior of the capsid was a surprise [14]. In 2016, we discovered that all known protruding features of spherical viruses were located on 21 gauge points in the Asymmetric Unit (AU) [14], see Figure 2. These gauge points were found by analyzing the distinct set of geometric locations determined specified by icosahedral point arrays. The strict adherence of all known spherical virus protrusions to these locations suggest that they likely play a critical structural role in the stability of viruses. Some viruses, such as the bacteriophage MS2 adhere so strongly to these constraints, that seemingly innocuous changes to their protruding features cause their virus structure to destabilize [23,24]. While point arrays dictate critical locations for viruses, they do not fully specify their structures.

After we have computed all the RMSD fits of point arrays which match the gauge points, we then decide which array(s) are the best fit [2]. In general, a better fitting point array has at least 0.5 Å lower RMSD, has at least one point per protein and more overall points of contact with the capsid proteins. When multiple arrays meet this criteria, they are listed as well. We will see that as viruses mature, they change the structure of point arrays with which they conform.

## 3. Results

We determined the best fitting point arrays [2] for 135 distinct viruses. We found 149 unique protrusions which determined the gauge points for these point array fits; which we classify as key structural protrusions. Furthermore, we want to understand why protruding features of viruses play such a critical role in the possible structural determination of the arrangements of proteins and even genetic material contained within the virus. There were 121 capsids which had a single best fit point array, and 14 which had two best fit point arrays, none of the capsids in our study had 3 or more best fit point arrays. Those capsids with two arrays, also had two different gauge points, leading to 14 more total protrusions than capsids studied. There are 38 dsDNA capsids, of which 34 have a single structural protrusion, 15 ssDNA capsids, of which 14 have a single structural protrusions, 17 dsRNA capsids, of which 14 have a single structural protrusion and 61 ssRNA capsids, with 55 having a single structural protrusion, see Table 1.

We found that there is no straightforward connection between T-number and Baltimore classification of spherical viruses, in agreement with Louten [18], however there are some interesting results to note. We found that nearly all ssDNA capsids (BC II) are T1 (87%), and most of ssRNA capsids (70%) are T3 or pT3. In addition, most of T3 capsids are only composed of ssRNA (87%), whereas T7, T16, and larger T-number capsids were found to only contain dsDNA. In summary, knowing the T-number provides limited information on the genome, as does only knowing the genome provide little information on the T-number. We only found two T9 structures, and three (pT169, T169), which may indicate a slight disagreement with the evolutionary representations suggested in [20], which stated that T9 should be well represented in viruses. We did not find any T12 or T19 capsids, though T27 and T169 were present.

### 3.1. Gauge Point vs Triangulation Number

We present the relationship between Triangulation number and gauge points in Table 2 and Figure 3. Overall T-number and gauge points are independent though a few points stick out. GP 2 and 5 are primarily used by T1 capsids, 9 of 13 protrusions and 8 of 11 protrusions, respectively. GP 15 (2-fold axes) is primarily used by T3 capsids with 7 of 8 protrusions. We also found that few viral protrusions are found between the 3 and 2 fold axes, GP 6 to 15, though GP 15 has 8 capsids. In general, knowing the location of a structural protrusion does not strongly indicate the T-number. Capsids with T-number greater than 16 all have a single structural protrusion located on GP 1 (5-fold) or GP 21.

**Table 2.**
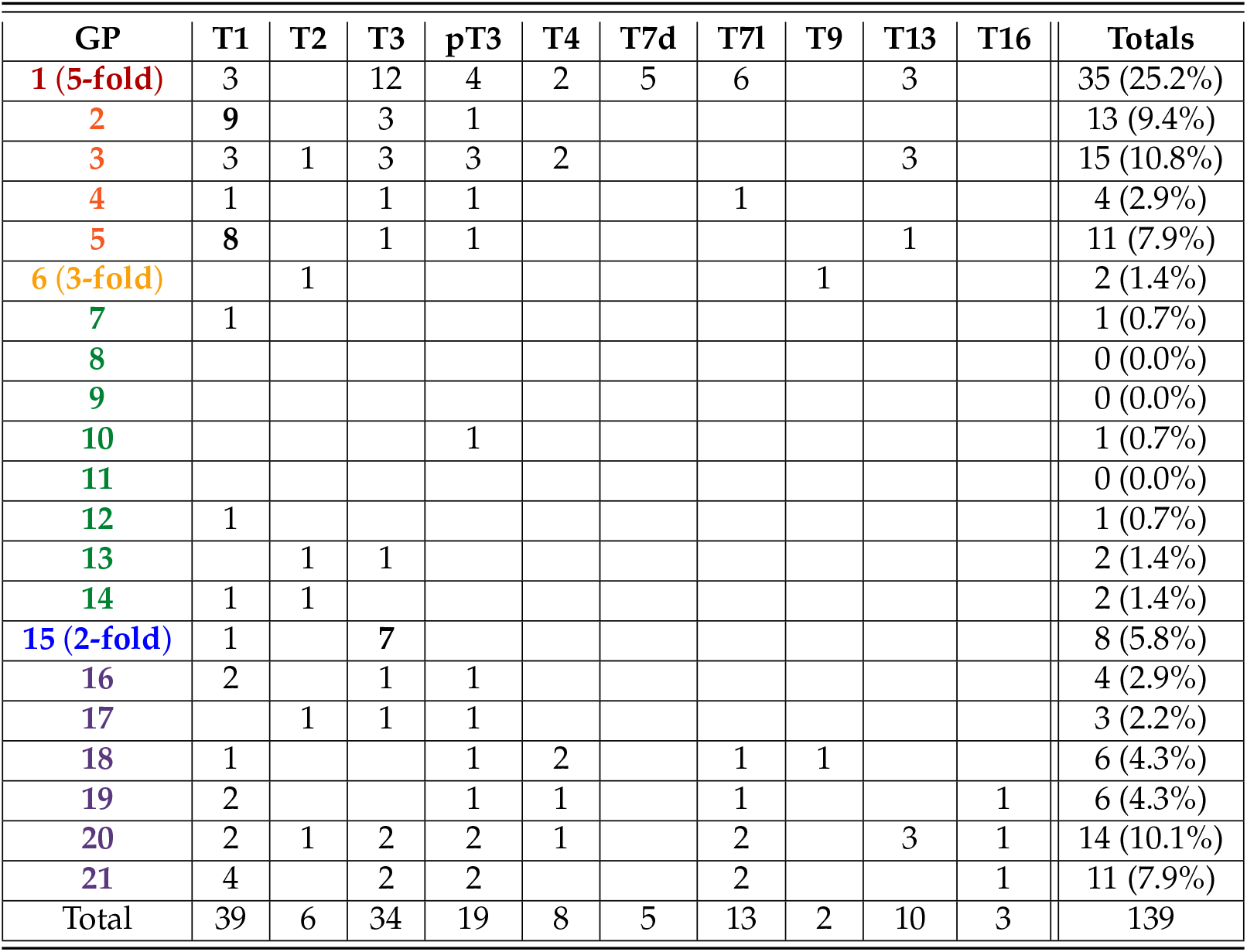
Gauge Points of Protruding Features vs T-number. In total we found 149 structurally significant protruding features within the AU when analyzing the 135 icosahedral capsids data set found in VIPERdb [1]. We found that most viruses have only one significant structural protrusion (90%) and the rest had two significant structural protrusions (10%); no virus has more than two. In general, knowing the location of a structural protrusion does not strongly indicate the T-number. However there are some exceptions, GP2 is mainly used by T1 capsids (69%), GP5 is mostly used by T1 capsids (73%) and the 2-fold axes (GP2) is primarily T3 (88%). We also found that few viral protrusions (6.5%) were found in the region from the 3-fold axes (GP6) to the 2-fold axes (G15), though the 2-fold (GP15) is well populated. There are 10 viruses with T-numbers larger than T16 not shown here, they all had a single structurally significant protrusion located at the 5-fold axes (GP1) or along the 5-2 region at GP21. Capsids with two structural protrusions ranged from from T1 to T13 and included each of the four genome types. This data can be visualized in Figure 3

**Figure 3.**
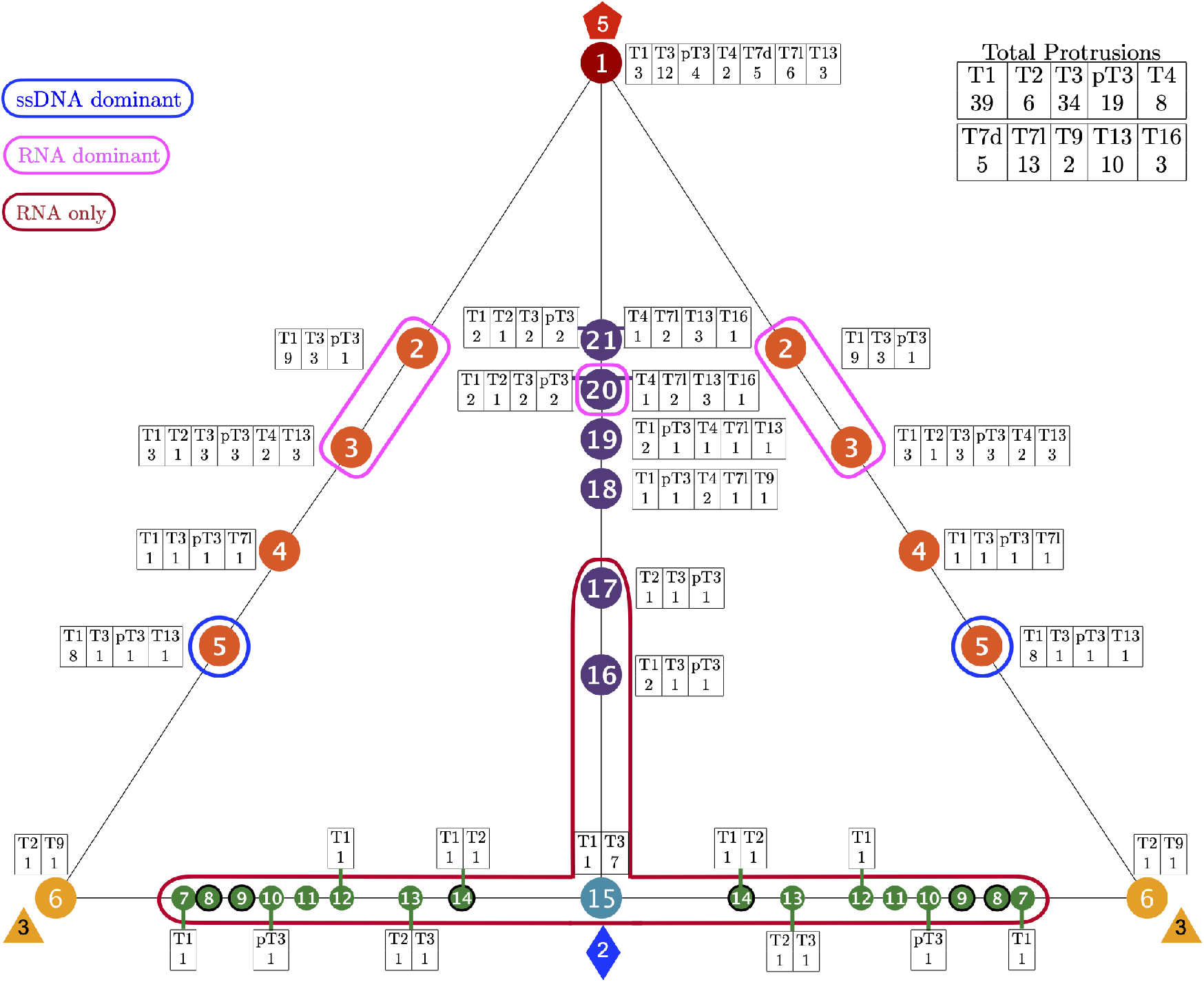
The 149 distinct structural protrusions and their corresponding T-numbers mapped onto the 21 unique gauge points of the Asymmetric unit (AU) [14]. The total number of protrusions for each T-number is shown in the upper right. The region encircled in red, GP 7 to GP 17 was found to only have RNA viruses with T-number 1 to 3. The regions encircles in pink (GP 2, 3 & 20) are dominated by RNA viruses (71% or more). Gauge point 5 is encircled in blue and primarily used by T1 capsid with ssDNA. The region between the 3-fold and 2-fold axes, GP 7 to 14 is not heavily used by any viruses. Overall T-number is not strongly predicted by gauge point location. There are 10 viral protrusions belonging to T21 and larger capsids not shown here. All of these capsids used GP 1 or GP 21. The angular locations of the gauge points are shown to scale on a flat AU face. There is a sizeable gap between GP 1 and GP 2 & GP 21, as there is between GP 5 and GP6.

### 3.2. Gauge Point vs Genome

We present the relationship between genomic composition and gauge points in Table 3 and Figure 4. Overall we find a strong relationship between the location of protrusions and genome composition. The most dramatic result was that only RNA viruses use Gauge Points 7 to 17, which we refer to as the upside down T/Up Tack (⊥) region, see Figure 3 and Figure 4. We also see that RNA capsids protrusions dominate GP2 (85%), GP3 (80.0%) and GP20 (71%). Capsids with ssDNA are also clearly dominant at GP5 (73%). There is also a clear relationship between dsDNA viruses between GP 1 and GP 21, representing 28 of 42 protrusions from 31 of the 38 dsDNA capsids. We also find that 13 of the 16 ssDNA protrusions are found between GP 1 (5-fold) and GP 5, accounting for 13 of the 15 ssDNA capsids, see Table 3 and Figure 4. Viruses with ssRNA compositions were spread all around the gauge points, with the largest number, 13 of 67 protrusions being found at GP 1. Similarly dsRNA capsids use a range of protruding feature locations. There were also some gauge points used by all genomes, GP 1, 2, 3 and 20, see Figure 4.

**Table 3.**
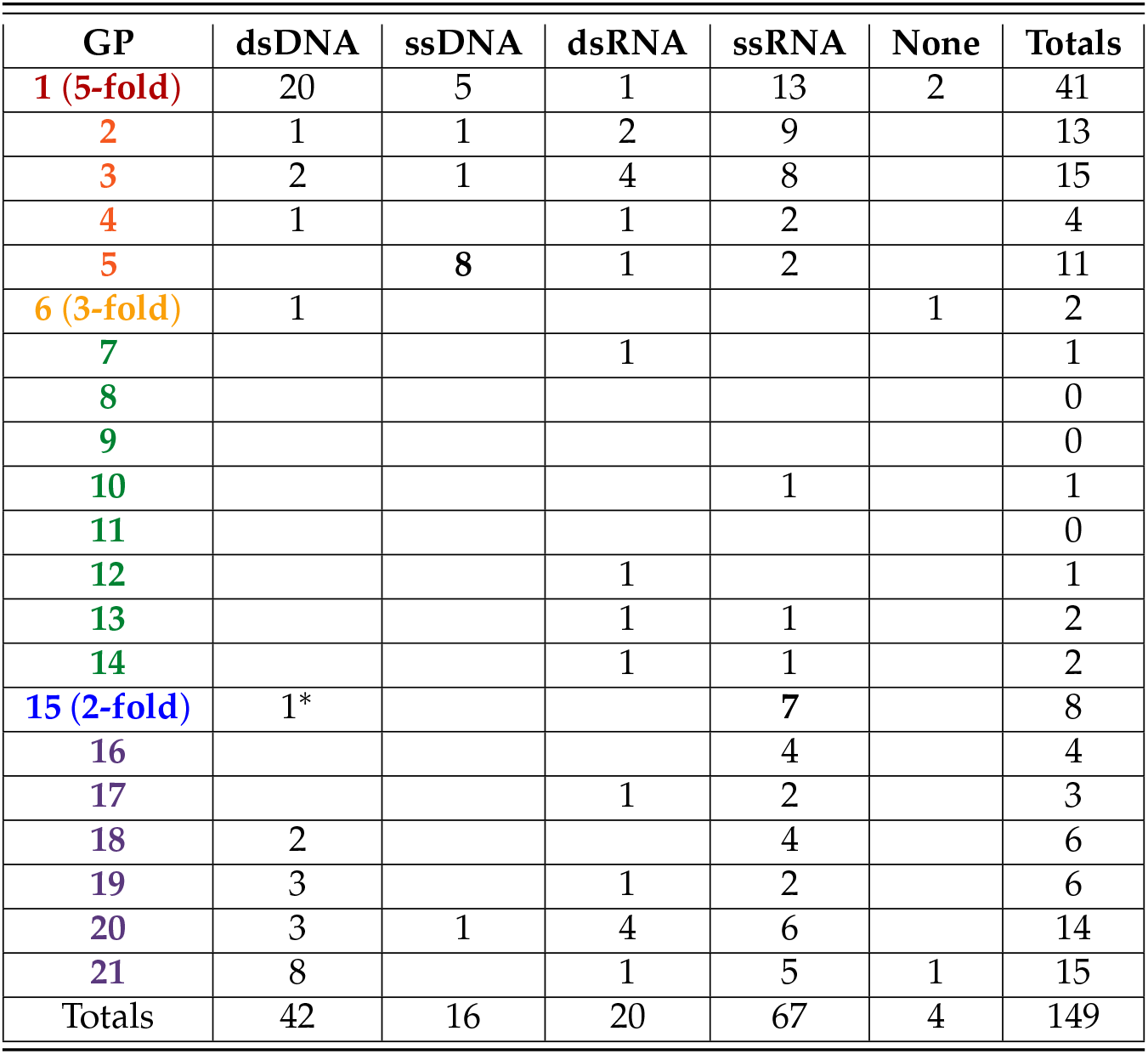
Gauge Points of Protruding Features vs Genome. The genomic compositions of the 149 structural protrusions of the 135 distinct capsids found in the VIPERdb [1]. We found that only RNA viruses are found to have protrusions from Gauge Point 7 to 17, which we refer to as the “T” region, see Figure 4. There is one notable exception, the aberrant T3 form of HBV, which is too small to actually encapsulate its DNA genome [25,26]. We also see that RNA capsids protrusions dominate GP2 (85%), GP3 (80%) and GP20 (71%). Capsids with ssDNA are also clearly dominant at GP5 (73%).

| GP | dsDNA | ssDNA | dsRNA | ssRNA | None | Totals |
| --- | --- | --- | --- | --- | --- | --- |
| **1 (5-fold)** | 20 | 5 | 1 | 13 | 2 | 41 |
| **2** | 1 | 1 | 2 | 9 |  | 13 |
| **3** | 2 | 1 | 4 | 8 |  | 15 |
| **4** | 1 |  | 1 | 2 |  | 4 |
| **5** |  | 8 | 1 | 2 |  | 11 |
| **6 (3-fold)** | 1 |  |  |  | 1 | 2 |
| **7** |  |  | 1 |  |  | 1 |
| **8** |  |  |  |  |  | 0 |
| **9** |  |  |  |  |  | 0 |
| **10** |  |  |  | 1 |  | 1 |
| **11** |  |  |  |  |  | 0 |
| **12** |  |  | 1 |  |  | 1 |
| **13** |  |  | 1 | 1 |  | 2 |
| **14** |  |  | 1 | 1 |  | 2 |
| **15 (2-fold)** | 1* |  |  | 7 |  | 8 |
| **16** |  |  |  | 4 |  | 4 |
| **17** |  |  | 1 | 2 |  | 3 |
| **18** | 2 |  |  | 4 |  | 6 |
| **19** | 3 |  | 1 | 2 |  | 6 |
| **20** | 3 | 1 | 4 | 6 |  | 14 |
| **21** | 8 |  | 1 | 5 | 1 | 15 |
| Totals | 42 | 16 | 20 | 67 | 4 | 149 |

**Figure 4.**
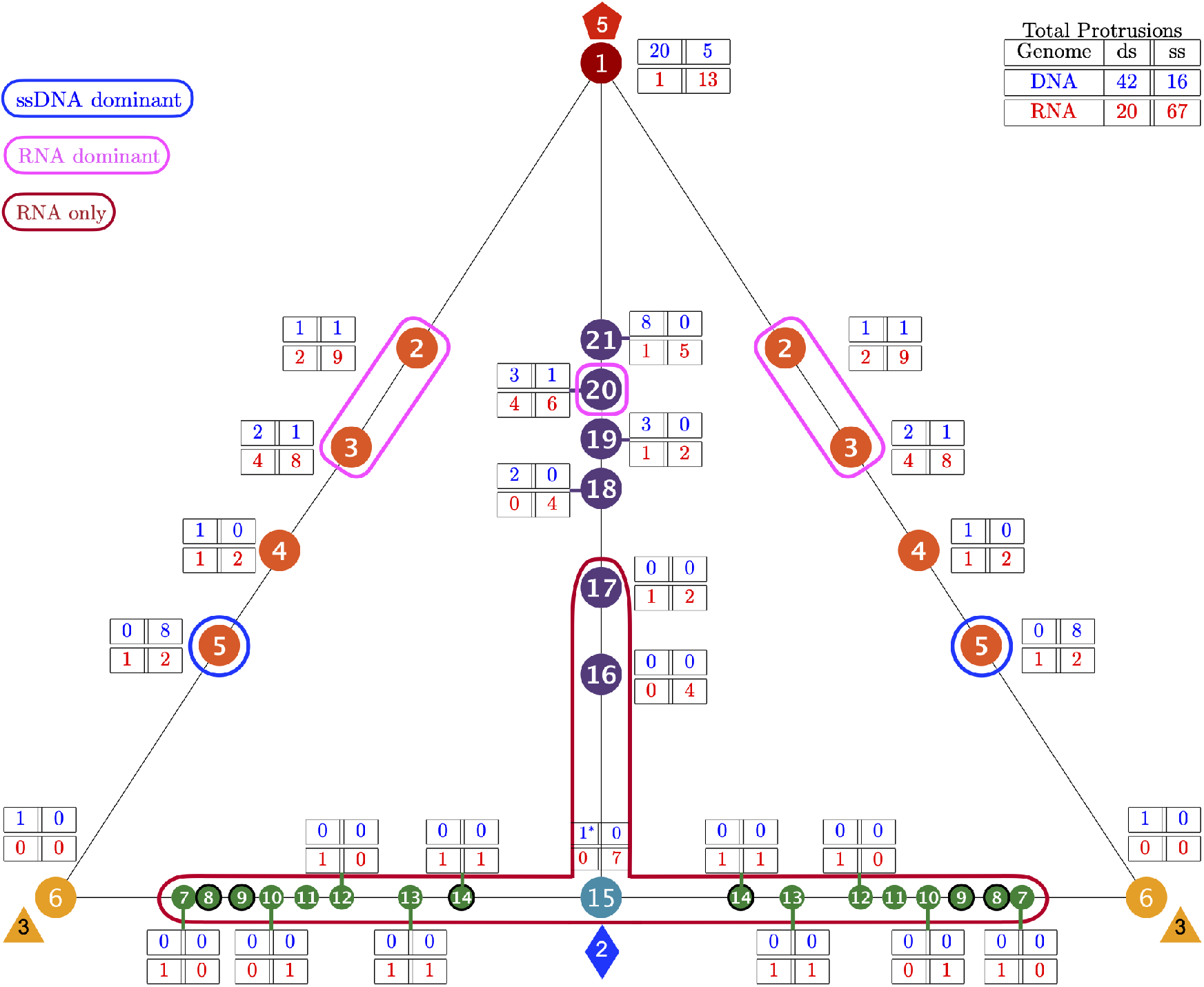
The 149 distinct structural protrusions and their corresponding genomes mapped onto the 21 unique gauge points of the Asymmetric unit (AU) [14]. The total number of protrusions for each genome is shown in the upper right. The region encircled in red, GP 7 to GP 17 was found to only have RNA viruses (Baltimore classifications III, IV, and V). There is one technical exception to this observation, HBV forms a mixture of T3 (5%) and a T4 (95%) capsids. The T3 capsid (6bvn) has two structural protrusions at GP 3 and GP 15, however the T3 capsid is too small to package the DNA genome inside [25,26]. The regions encircles in pink (GP 2,3 & 20) are dominated by RNA viruses (71% or more). Gauge point 5 is encircled in blue and primarily used by T1 capsid with ssDNA (73%). Gauge points 1, 4, 18, 19 and 21 are equally utilized by RNA and DNA viruses. This visualization elucidates the new utility of Viral Phrenology, as a predictor of genome composition based on location of protrusions.

### 3.3. Gauge Points 1 to 6

Gauge point 1 is the 5-fold axis and is used by DNA and RNA viruses. Most dsDNA viruses use GP 1 (20 of 42 capsids), accounting for 20 of its 38 dsDNA protrusions located here. We found 13 capsids which were T16 and larger were only composed of dsDNA. There are 10 capsids not shown in Table 2, pT21(2), pT25(2), PT27, T28d, pT31, T43, pT169, T169, which all used Gauge Point 1 or 21. Moving from the 5-fold axes along the great circle to the 3-fold axes (GP 2 to GP 6), we find that GP 2 is dominated by RNA, 11 of 13 viruses (84.6%). We find that GP 3 is also dominated by RNA, 12 of 16 viruses (75.0%). We find that GP 4 was barely used, of the four viruses that did, two of them also had protrusions elsewhere. Gauge point 5 was dominated by ssDNA, 8 of 11 viruses (72.7%), see Figure 5 and Figure 6. Lastly Gauge Point 6 seems to be largely unused by icosahedral capsids. While some Adenoviruses are known to have trimeric bundles on the 3-fold axes, we did not have access to these structures arranged with icosahedral symmetry. Curiously, the best fit protrusions at the 3-fold axis was a bacterial micro-compartment [27], which does not resemble any other capsids we studied. We postulate that there could be an interaction with 3-fold protrusions and viruses that is being avoided. The locations of the protein protrusions near the 3-fold axes also allow for gauge point 5 in principle, however none of the capsids which correspond to point were a good fit due to point arrays penetrating protein volumes. This indicates there is something substantially different about this structure and other viral capsids. The only virus we found with a structurally significant 3-fold protrusion was a member of the Reoviridae family [28], which also had a protrusion at GP 3. The best fitting point array (GP 3) had an RMSD of 2.1 Å, which was 0.5Å better than the 3-fold array with RMSD of 2.6Å, and by our prior classification schemes [2] would be considered the best array, however we have included it here due to it being unusual. The only result that used gauge point 6 was the bacteria compartment 6mzx [27].

**Figure 5.**
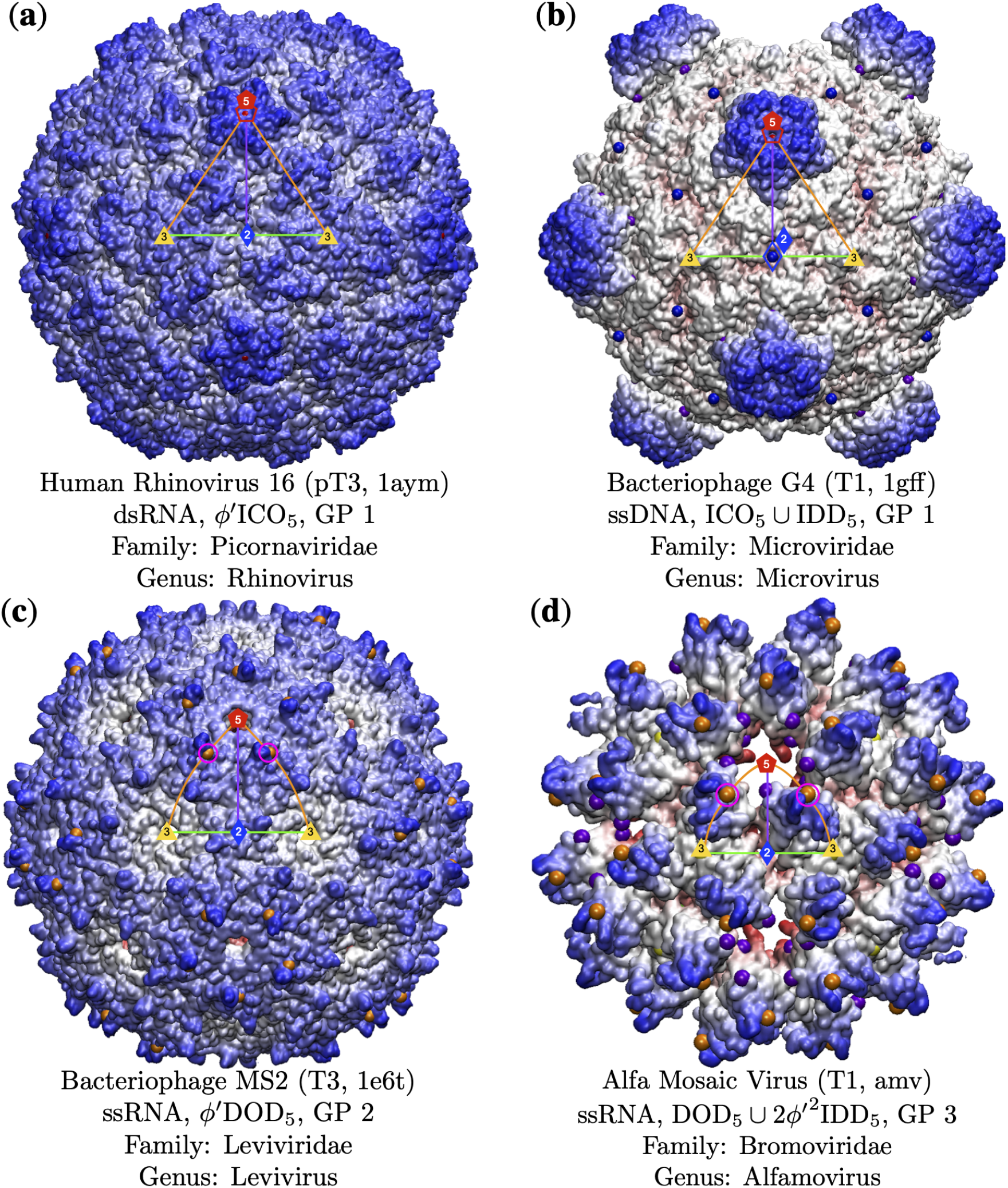
Examples of capsids with Gauge Points (GP) 1 to 3. The gauge points which correspond to the key structural protrusions are shown in pink circles. The Triangulation number, pdb id, genome composition, point array classification [2]. (**a**) Human Rhinovirus [29] has a structural protrusion on the 5-fold axes (GP 1). There is another protrusion along the 5-3 GC, however none of the point arrays corresponding to these protrusions fit the virus capsid well [2]. (**b**) Bacteriophage G4 [30] uses GP 1 for its protrusions. GP 21 was also a possible fit, however none of the point arrays associated with this gauge point were a good fit for the capsid. (**c**) Bacteriophage MS2 [31] has surface loops at GP 2, as shown. Previous experimental work showed that while these loops do not appear structural in nature, changes to the amino acids adjacent to GP 2 was nearly impossible [2,23,24]. As all of the proteins in MS2 are chemically identical, this restriction to modification extends to all protrusions of MS2 pre-assembly. (**d**) Alfa Mosaic Virus [32] uses GP 3 for its protrusions and has remarkable agreement (6 radial levels,[2] with the double point array DOD_5_ ~ 2*ϕ*′IDD_5_ and with an RMSD of 1.6Å.

**Figure 6.**
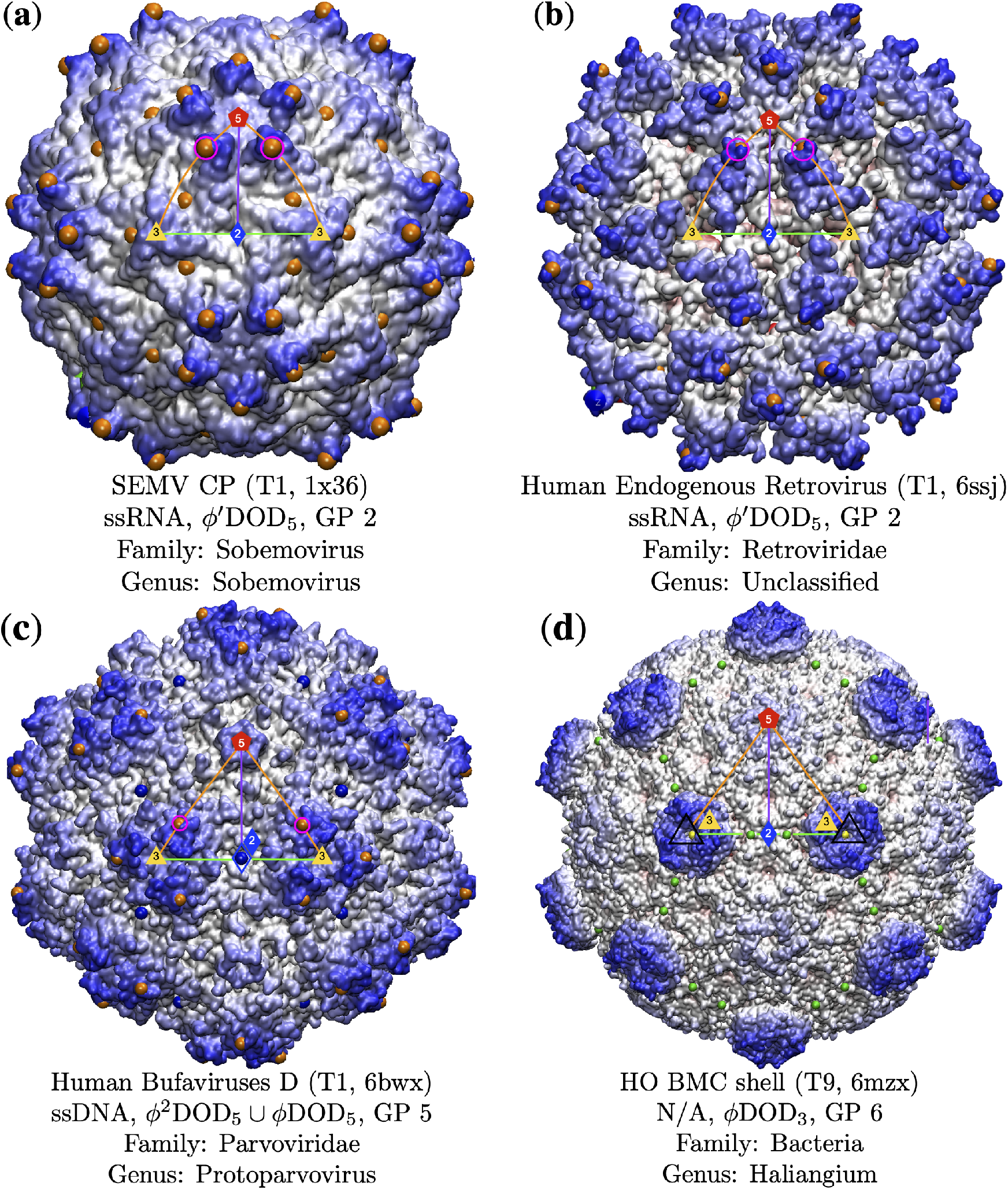
Gauge points along the 5-3 arc (GP 1 to 6). Gauge points which correspond to the key structural protrusions are encircled in pink. Gauge Points 2 & 3 are dominated by RNA capsid protrusions (84.6% and 80.0%, respectively). (**a**) Sesbania Mosaic Virus [33] is GP 2 but it also has protrusions near the 3-fold axes, however these locations are over 5Å below the ones at GP2, and do not correspond to any point arrays that fit the capsid. We can see that despite using the same gauge point, these two capsids look quite distinct. (**b**) Human Endogenous Retrovirus [34] also uses GP 2, though it looks quite distinct from SEMV, with relatively large openings along the 3-2-3 arc. (**c**) Human Bufaviruses D [35] uses GP 5 for its protrusion. Note the valley between proteins resulting in a lower surface along the 3-2-3 arc. (**d**) An icosahedral bacterial micro-compartment [27] which uses GP 6, only one of two structures in our library to do so. To the eye this structure is clearly distinct from all others in this paper and we speculate that it is advantageous for viruses to not present as bacterial micro-compartments.

#### 3.3.1. T7d Viral Capsids

We initially had difficulty finding the best fitting point arrays for the T7d Papillomaviridae (5kep[36], 5keq[36], 3iyj[37]) and Polyomaviridae (1sva[38], 6gg0[39]) capsid families [14], which includes SV40. Each of these capsids are composed entirely of pentamers, resulting in 360 capsid proteins instead of the expected 420 (T7). However, each of these capsids have at least one viral protein found at the 5-fold axes, inside or beneath the pentamers located on 5-fold axes, though this protein is not arranged with global icosahedral symmetry [38,40]. The coordinates for these proteins are not found in their pdb coordinates. However when we take these into account, each of these 5 capsids are GP1. It is possible that Bovine Papillomavirus (3iyj [37]) and BK Polyomavirus Like Particle (6gg0 [39]) could turn out to have structural protrusions at GP 4, as some of the best fitting point arrays we found were located here, however none of these point arrays made contact with all the capsid proteins. It is possible that including more coordinate data from the genomes could offer additional point array fits. For the analysis of 6gg0, we fit this capsid neglecting the neutralizing antibody, though we plan to investigate the induced conformation changes in future work.

#### 3.3.2. Parvoviradae Family

The Parvoviridae Family are non-enveloped T1 capsids with ssDNA that infect a wide range of hosts from vertebrates and invertebrates [41] and all use Gauge Point 5, making up 8 of the 11 viruses at this location. In Figure 7 you can see that while the surfaces vary, they all use the same locations for their protrusions. One fascinating member of this family is M. spretus, a VLP [42] that was discovered as an endogenous viral elements incorporated in Mus spretus, an Algerian mouse. It’s genetic similarity to other parvoviruses indicate that its less than 2 million years old. We note that this reconstituted capsid still uses GP 5, indicated this location is likely of great importance to parvoviruses, perhaps critical to the family overall. The H-1 parvovirus capsid has biomedical applications, and a genome-free VLP has been developed as an antitumor gene delivery vector [43]. Not all members of this family have great point arrays fits, as in the case of Adeno-Associated Virus (AAV2). These protrusions are slightly shifted off GP 5, so that the gauge point rests at the base of them. However, this is still the best fit array considering all other arrays available. The AAV2 structure was determined by cryoEM using several advancements in techniques, including per-particle CTF refinement and corrections for Ewald sphere curvature, to improve resolution [44]. We intend to study this and related to structures in the future to determine the source of the deviation of the gauge point from the protrusions. We also examined Adeno-Associaed Virus in-complex with its cell receptor AAVR [45], see Figure 7. Here the complexed structure still used GP 5, however we found that the internal arrangement of constraints shifted outer point arrays from *ϕ*′^2^ICO_3_ to *ϕ*′^2^DOD_5_ which is the sister array at GP 5 with one interior radial level difference [2]. These additional three interior point arrays represent other internal structure changes allowed by this complexed structure, which we therefore expect this complex to be more stable.

**Figure 7.**
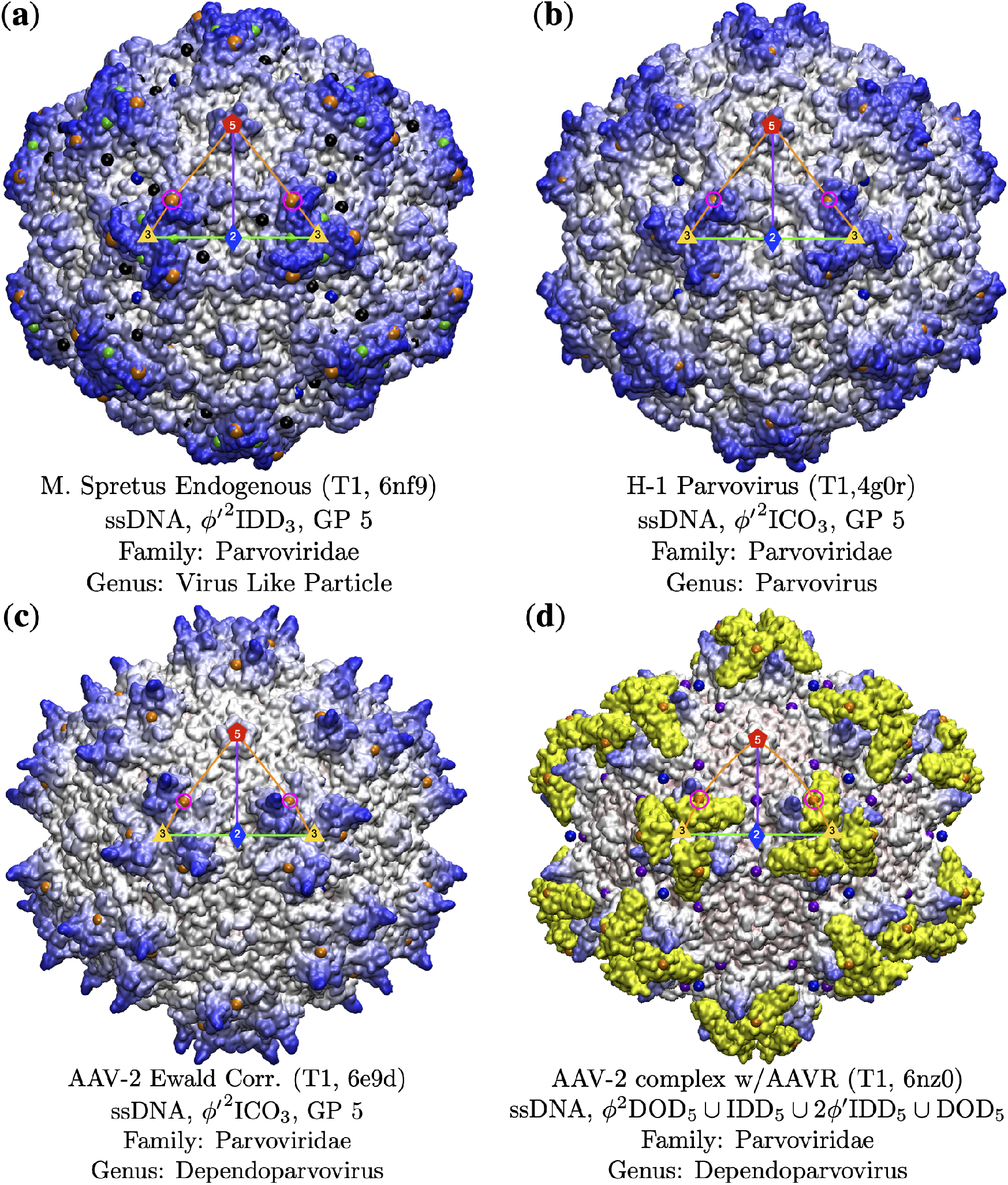
The Parvoviridae Family are non-enveloped T1 capsids with ssDNA that infect a wide range of hosts from vertebrates and invertebrates [41] and all use Gauge Point 5, making up 8 of the 11 viruses at this location. As you can see in these images, their surfaces vary, however they all use the exact same gauge point 5 for their structural protrusions. (**a**) M. spretus is a VLP which was discovered as an endogenous viral element. (**b**) The H-1 parvovirus capsid has biomedical applications [43]. (**c**) Adeno-Associated Virus (AAV2) has protrusions that are slightly shifted off GP 5 and was found using cryoEM[44]. No other gauge points and associated point arrays fit this structure. (**d**) Adeno-Associaed Virus in-complex with its cell receptor AAVR, shown in yellow [45]. This combined structure still used GP 5, however we see that the internal arrangement of constraints has shifted outer point arrays from *ϕ*′^2^ICO_3_ to *ϕ*′^2^DOD_5_ which is the sister array at GP 5 with one interior radial level difference [2].

### 3.4. Gauge Points 7 to 17: RNA viruses only

There are few virus capsids with protrusions along the 3-2-3 arc of the AU, GP 7 to 15, see Figure 3 and Figure 4. Most of the viruses in this region use the 2-fold axes for their protrusions. It is possible that having protrusions in this region lead to smaller capsid volumes, making it more difficulty for DNA packing. Overall we found that GP 7 to 17 are only RNA viruses (BC IV & BC V), accounting for 28% of all RNA viruses in our library (22 of 29). Gauge point 7 to 14 is sparsely populated, with 7 of 79 RNA viruses. Gauge point 15 to 17 had 15 viruses. The only exception was the T3 HBV capsid, which has a protrusion at the 2-fold axis (GP 15), however this form of the capsid is unable to fit the DNA genome inside. It is interesting to note that at this stage of assembly, HBV contains pre-genomic RNA [25,26]. See Figure 8 and Figure 9 for examples of viruses along GP 7 to 17. We also note that no viruses were found that use GP 8, 9 or 11.

**Figure 8.**
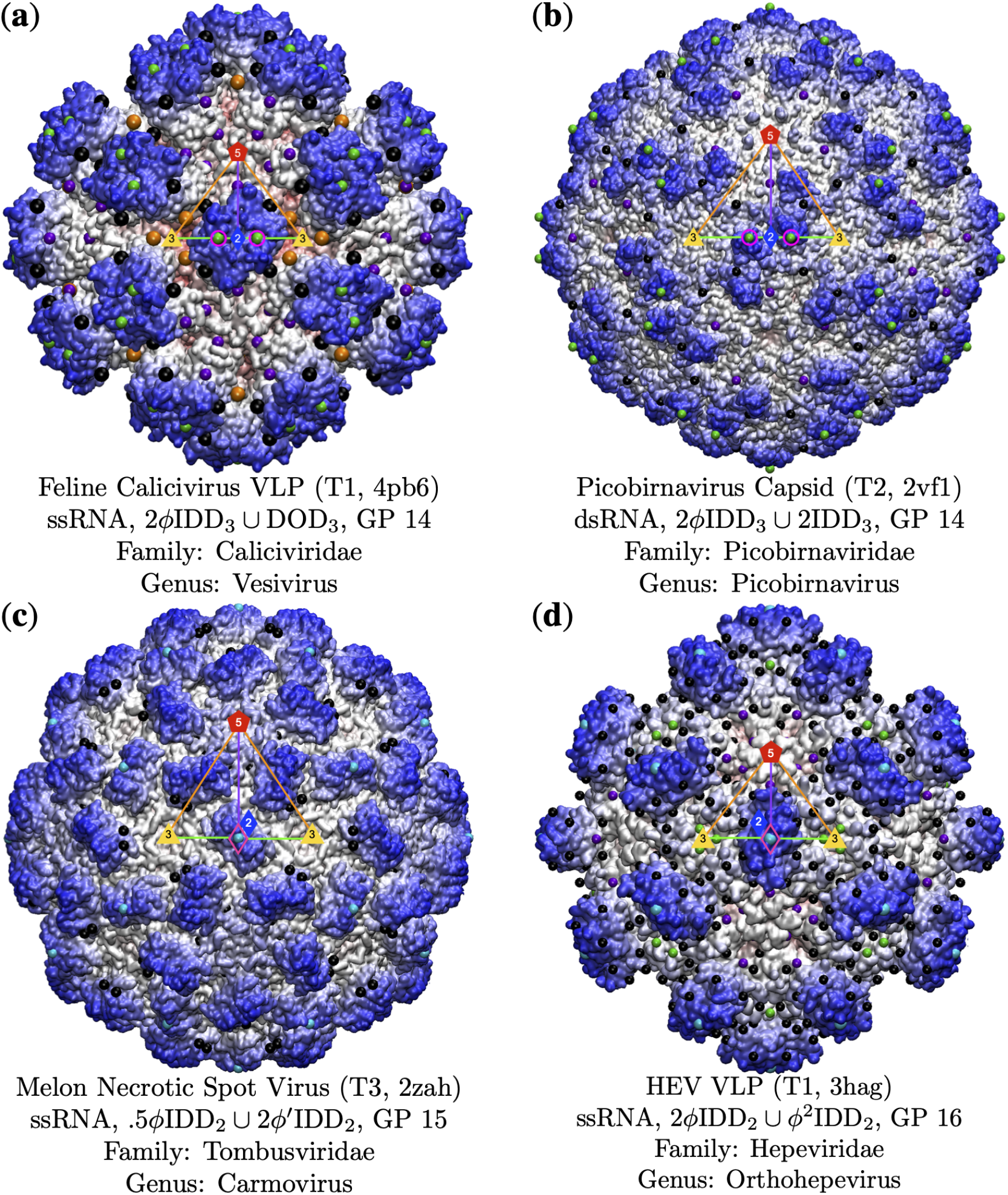
Gauge Points 7 to 17 are exclusively RNA viruses. The capsid surfaces are diverse and their protrusions are well located on the gauge points. (**a**) Feline Calicivirus [46] is a VLP with protrusions that flair up near GP 14. (**b**) Picorbirnavirus is a T2 capsid [47] which also uses GP 14, though its surface differs considerably from Feline Calicivirus. (**c**) Melon Necrotic Spot Virus [48] has a structural protrusion at GP 15. There is another protrusion at GP 3, however none of the associate point arrays fit the capsid well as well. (**d**) Hepatitis E VLP [49] has excellent agreement with its point arrays [2] and uses the 2-fold axes.

**Figure 9.**
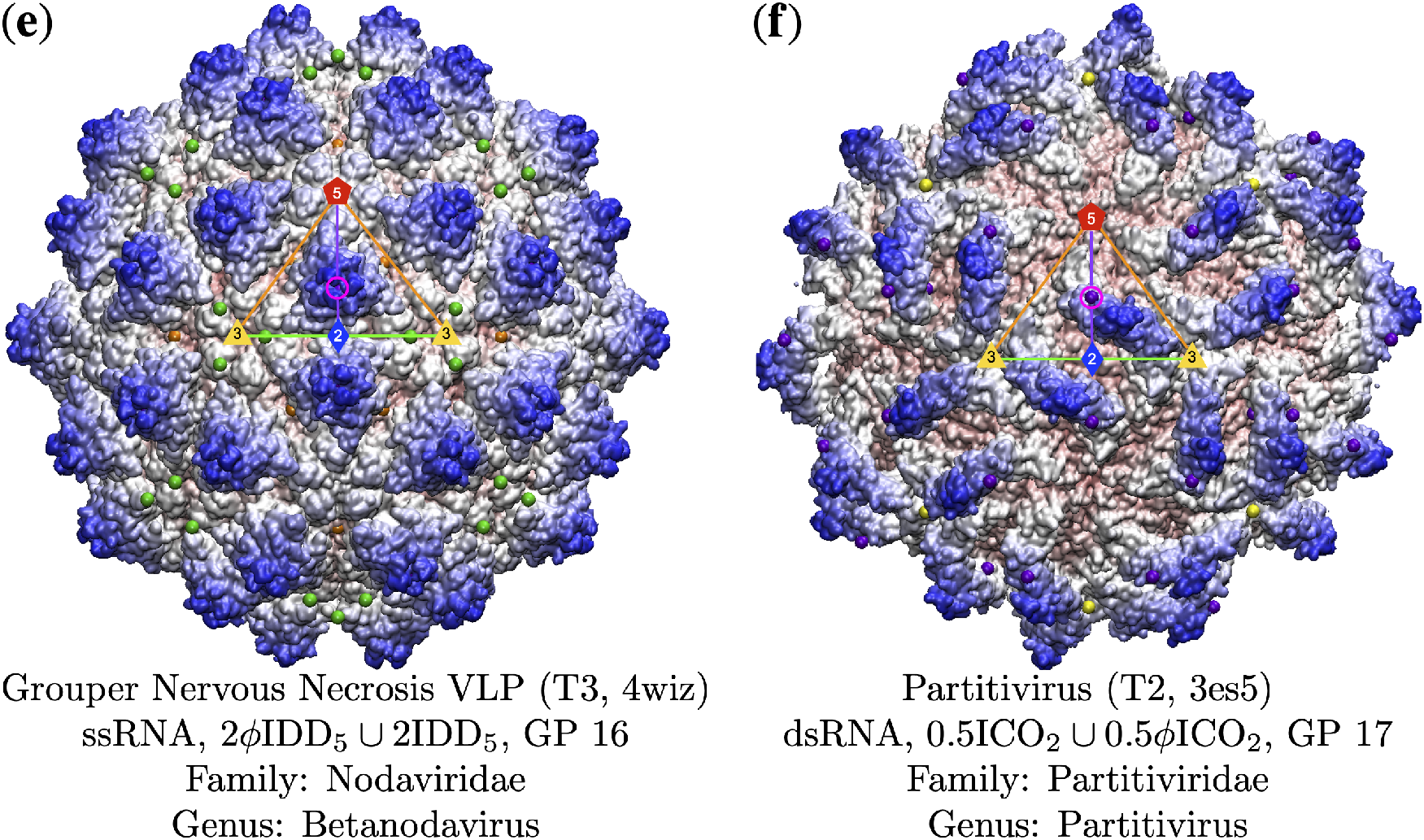
Gauge Points 7 to 17 continued. This region is home exclusively to RNA viruses. (**e**) Grouper Nervous Necrosis VLP [50] uses GP16 that is slightly embedded with the protrusion. Overall this capsid does not have a great fit with the point arrays, as can be seen with the floating green points. (**f**) Partitivirus [51] has an unusual protrusion that connects to the viral surface in the bulk region, however the edge of it intersects the 5-2 arc at GP 17.

### 3.5. Gauge Points 18 to 21

We now turn our attention to the rest of the arc between the 5- and 2-fold axes, see Figure 10. Gauge point 18 and 19 are sparsely populated, with 6 viruses each and a mixture of RNA and DNA composition, see Table 3, Figure 3, and Figure 4. Gauge point 20 is dominated by RNA viruses, 10 of 14 capsids. Finally GP 21 is equally used by DNA and RNA viruses. This is also the second largest location for dsDNA capsids with 8 of 42 being found here, in-total GP 1 and GP 21, account for 28 of 42 dsDNA capsids. There are also 4 viruses with T21 or larger architecture that use GP 21, see Table 2.

**Figure 10.**
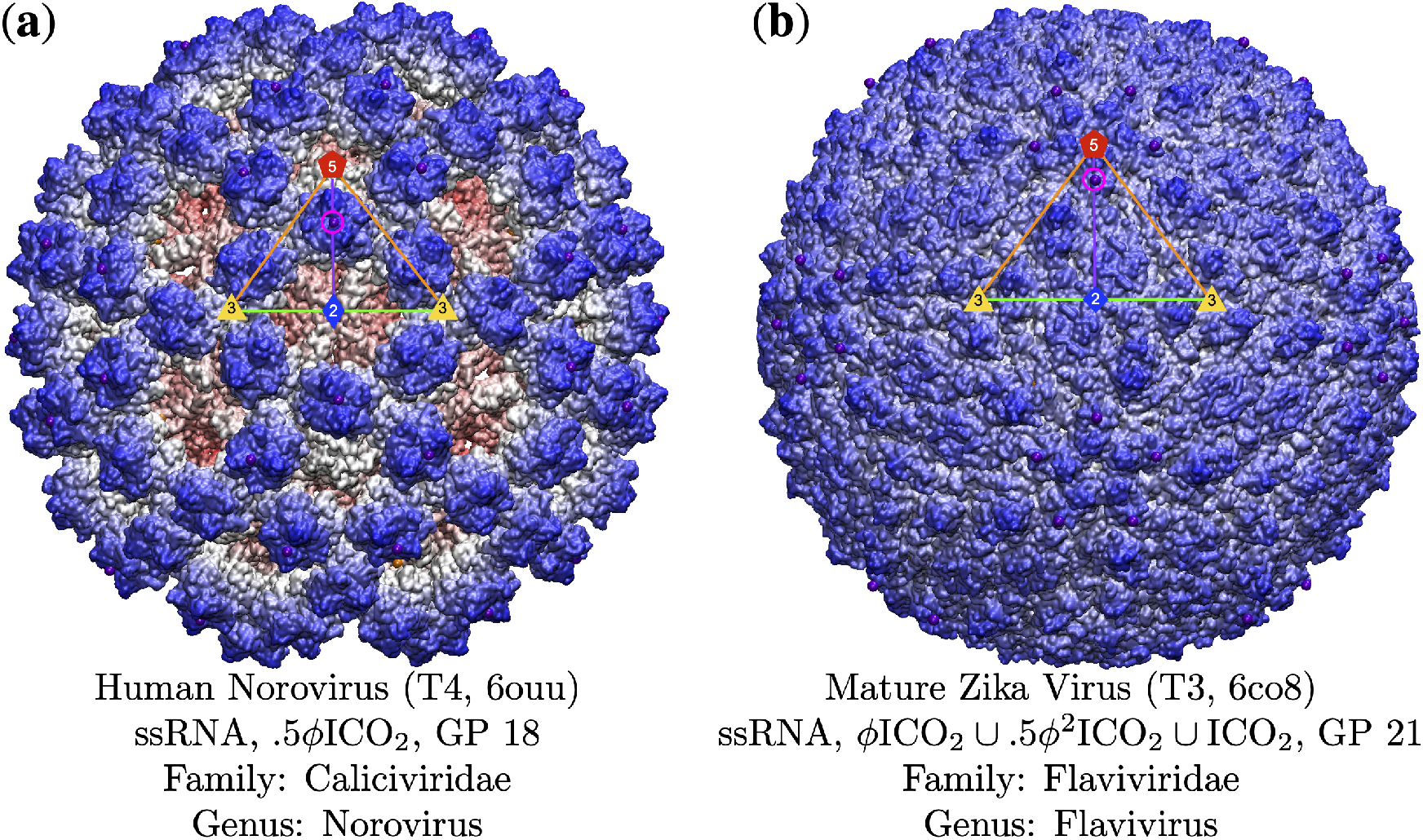
Gauge points 18 to 21. (**a**) Human Norovirus [52] is a T4 capsid with two protruding features, however only the protrusion at GP 18 has an associated point array that fits the capsid well. (**b**) The Mature Zika Virus [53] has the some of the least differentiated protrusions out of all the viruses in our database. We initially identified protrusions at GP 1, 3, 8, 10, and 21. However, only the point arrays with Gauge Point 21 shown above had low RMSD scores (1.6Å with GP 21 arrays vs. 3.2Å with GP 8 arrays).

## 4. Conclusions

Our work shows that Viral Phrenology, the study of spherical capsid protrusions to deduce the genomic composition of a virus, is a useful tool in understanding the structure and assembly of viruses. While the importance of protrusions to infection has been long established, the patterns and rules they obey is just now being revealed. We show that gauge points and their associated point arrays are an important compliment to the traditional Triangulation number and Baltimore classification of viruses. The implications and utility of this work are broad, especially when considering potential modifications of viruses and designing new virus-like particles (VLPs). While the reason for viruses adhering to point arrays are still poorly understood, the rules and patterns which viruses clearly follow are becoming more apparent.

Point arrays continue to be an important tool for understanding viruses, from knowing where it is permissible or not to substitute or modify surface features, to helping explain differences in vibrational properties of chemically identical proteins, to understanding binding affinity on surfaces [2]. Now we see a clear connection to genomic composition as well, which should be considered when modifying viral surfaces and designing new VLPs. The origins of these restrictions is not yet clear, it could very well be evolutionarily fixed, as is possible in the Parvoviradae or Papillomaviridae families.

It is speculated that dsRNA viruses evolved from positive-sense single-strand RNA viruses which then lead to negative-sense single-strand RNA viruses [54]. This could account for each genomic type using all of the gauge points in similar numbers. There is also reason to believe that protrusions along the 2-3-2 section of the AU will lead to smaller capsid volumes than DNA capsids could use, though more study is needed.

It is now clear that point arrays, Baltimore classification and Triangulation number are all communicating different information about the virus structures. It is our belief that understanding point arrays will provide a connection between Triangulation number and Baltimore classification, as an understanding of the gauge points has lead to an understanding of genomic composition. It is also clear that some T-numbers have a strong connection to certain gauge points, which then limit the potential point array structures.

## Author Contributions

“Conceptualization, D.R. and D.W.; Construction of virus library, D.R.; initial classifications, D.R. validations, D.W.,; formal analysis, D.W.; investigation, D.R.; resources, D.W.; data curation, D.R.; writing—original draft preparation, D.W.; writing—review and editing, D.W. and D.R., supervision, D.W.; All authors have read and agreed to the published version of the manuscript.”.

## Data Availability Statement

Our pdb library list is available in the Appendix. Any other data is available upon request.

## Acknowledgments

D. Wilson would like to thank Jan Tobochnik for his assistance with editing and Vijay Reddy for his tireless answering of questions about VIPERdb.

## Abbreviations

The following abbreviations are used in this manuscript:

AU: Asymmetric Unit
BC: Baltimore Classification GP Gauge Point
PA: Point Array
TX: Triangulation number X, *e.g*. T3 or T7.

## Appendix A Data Set

From the over 600 viral capsids deposited in VIPERdb [1], we found 135 distinct capsids with unique families, genera, Triangulation numbers and lowest resolutions. When there was a tie for lowest resolution, both structures were used. Resolution ties accounted for 6 additional capsids.

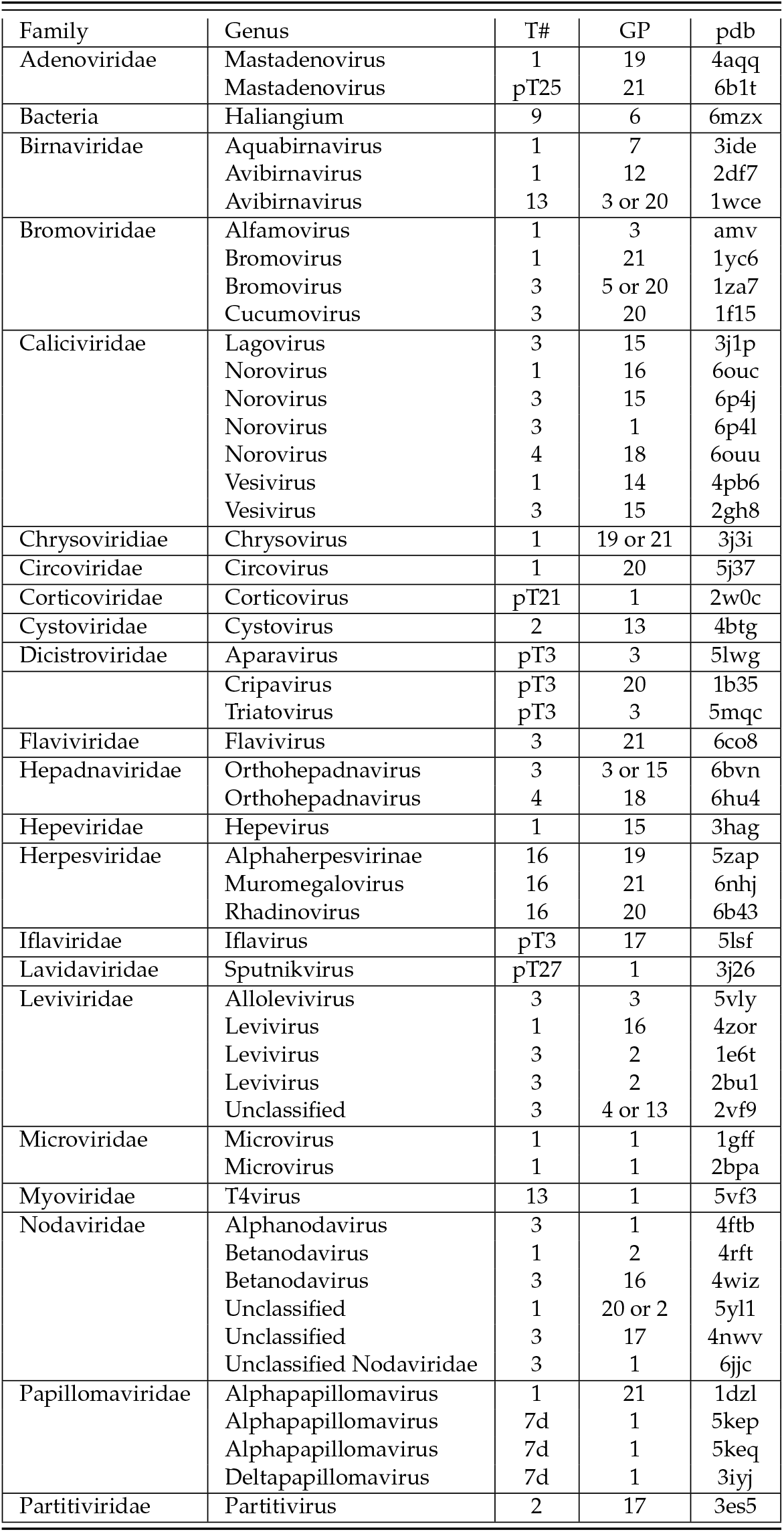
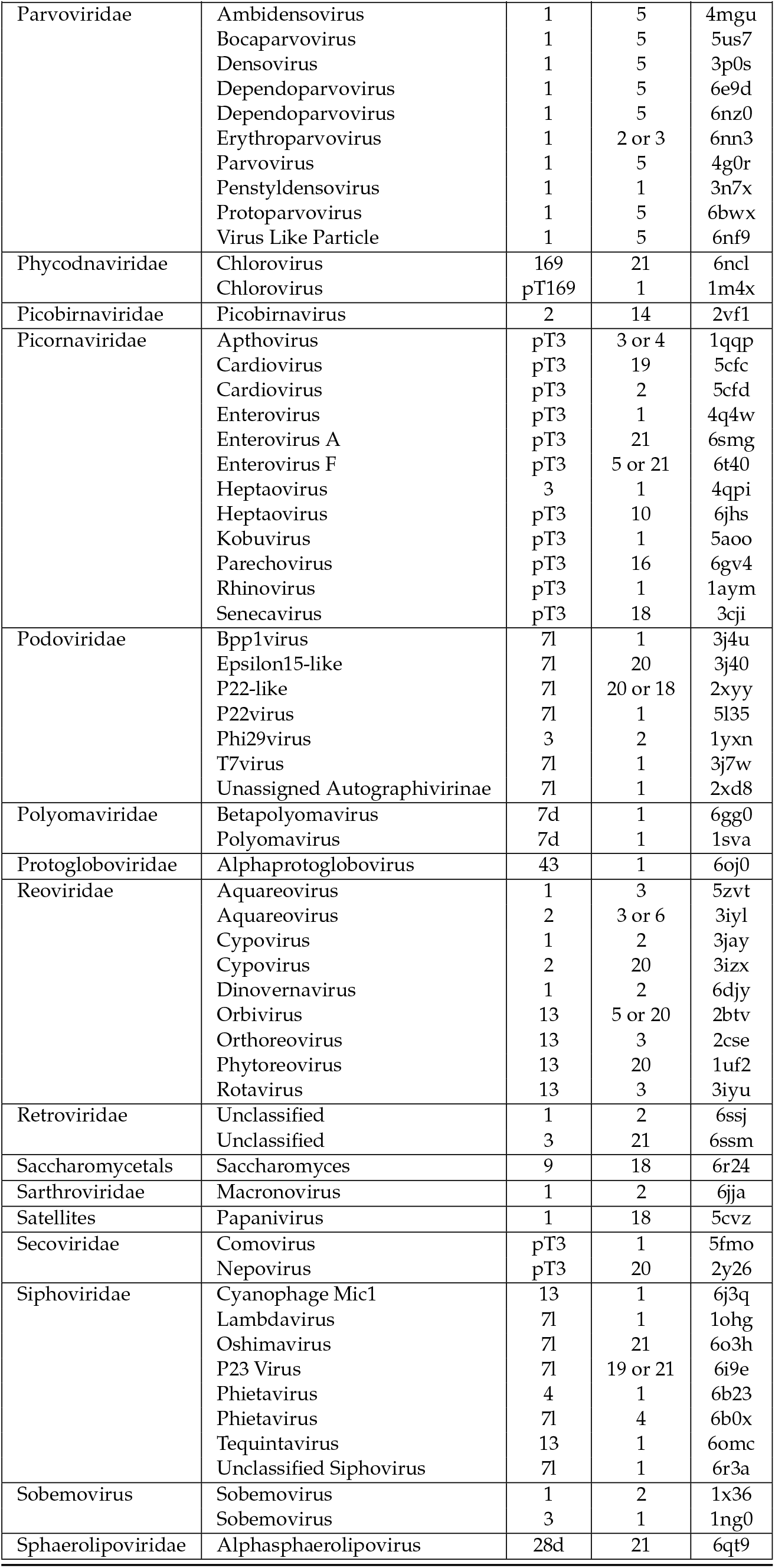
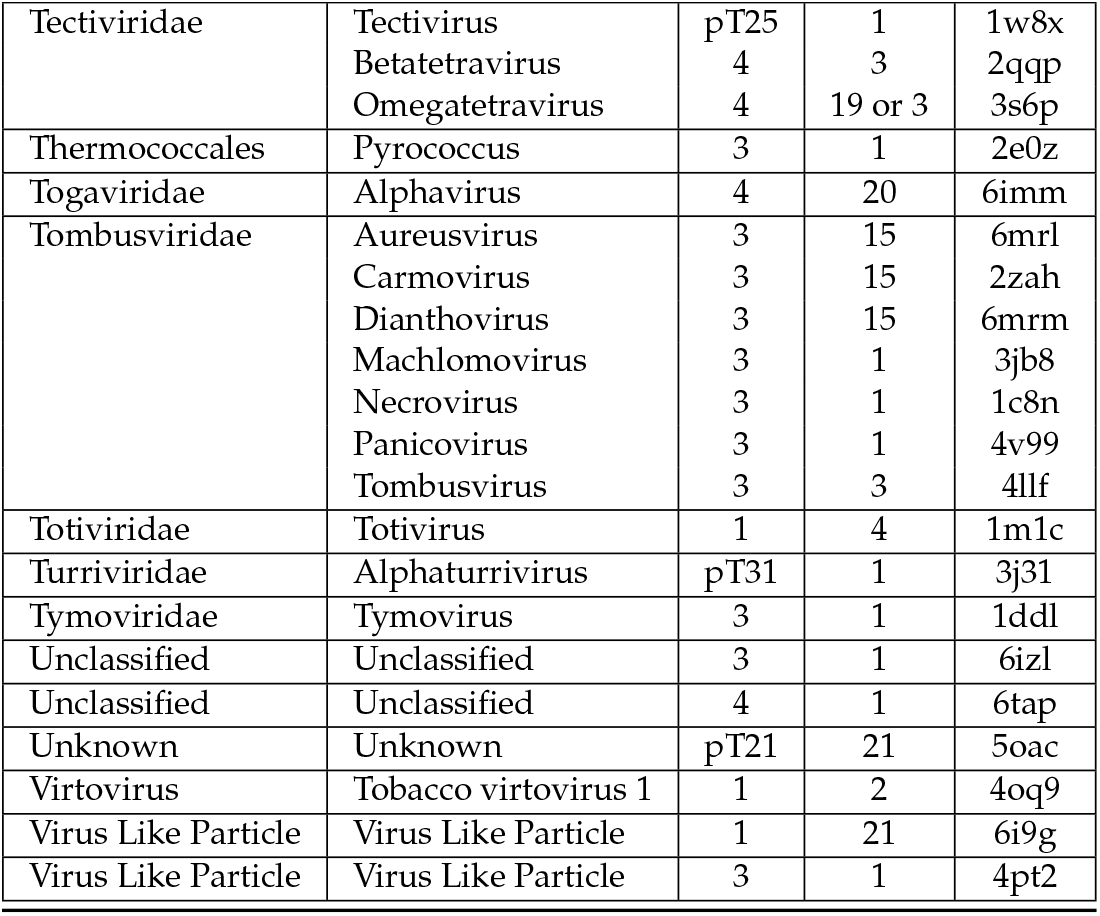

## REFERENCES

1. Montiel-Garcia, D.; Santoyo-Rivera, N.; Ho, P.; Carrillo-Tripp, M.; Brooks, C.L.; Johnson, J.E.; Reddy, V.S. VIPERdb v3.0: A structure-based data analytics platform for viral capsids. Nucleic Acids Research 2021, 49, D809–D816. doi:10.1093/nar/gkaa1096.

2. Wilson, D.P. Unveiling the hidden rules of spherical viruses using point arrays. Viruses 2020, 12, 1–33. doi:10.3390/v12040467.

3. Keef, T.; Twarock, R. Affine Extensions of the Icosahedral Group with Applications to the Three-dimensional Organisation of Simple Viruses. Journal of Mathematical Biology 2009, 59, 287–313. doi:10.1007/s00285-008-0228-5.

4. Keef, T.; Wardman, J.P.; Ranson, N.A.; Stockley, P.G.; Twarock, R. Structural constraints on the three-dimensional geometry of simple viruses: case studies of a new predictive tool. Acta Crystallographica Section A Foundations of Crystallography 2013, 69, 140–150. doi:10.1107/S0108767312047150.

5. Janner, A. Strongly correlated structure of axial-symmetric proteins. II. Pentagonal, heptagonal, octagonal, nonagonal and ondecagonal symmetries. Acta Crystallogr D Biol Crystallogr. 2005, 61, 256–68.

6. Janner, A. Crystallographic structural organization of human rhinovirus serotype 16, 14, 3, 2 and 1A. Acta Crystallogr A 2006, 62, 270–286.

7. Janner, A. Form, Symmetry and Packing of Biomacro-molecules. I. Concepts and Tutorial Examples. Acta Crystallographica Section A: Foundations of Crystallography 2010, 66, 301–311. doi:10.1107/S0108767310001674.

8. Janner, A. Form, symmetry and packing of biomacromolecules. II. Serotypes of human rhinovirus. Acta Crystallographica Section A Foundations of Crystallography 2010, 66, 312–326. doi:10.1107/S0108767310001698.

9. Janner, A. Form, Symmetry and Packing of Biomacromolecules. III. Antigenic, Receptor and Contact Binding Sites in Picornaviruses. Acta Crystallographica Section A: Foundations of Crystallography 2011, 67, 174–189. doi:10.1107/S0108767310053584.

10. Janner, A. Form, Symmetry and Packing of Biomacromolecules. IV. Filled Capsids of Cowpea, Tobacco, MS2 and Pariacoto RNA Viruses. Acta Crystallographica Section A: Foundations of Crystallography 2011, 67, 517–520. doi:10.1107/S0108767311035513.

11. Janner, A. Form, Symmetry and Packing of Biomacromolecules. V. Shells with Boundaries at anti-nodes of Resonant Vibrations in Icosahedral RNA Viruses. Acta Crystallographica Section A: Foundations of Crystallography 2011, 67, 521–532. doi:10.1107/S010876731103577X.

12. Zappa, E.; Dykeman, E.C.; Twarock, R. On the Subgroup Structure of the Hyperoctahedral Group in Six Dimensions. Acta Crystallographica Section A: Foundations and Advances 2014, 70, 417–428, [1402.3136v2]. doi:10.1107/S2053273314007712.

13. Zappa, E.; Dykeman, E.C.; Geraets, J.A.; Twarock, R. A Group Theoretical Approach to Structural Transitions of Icosahedral Quasicrystal s and Point Arrays. Journal of Physics A: Mathematical and Theoretical 2016, 49, 175–203, [1512.02101v2]. doi:10.1088/1751-8113/49/17/175203.

14. Wilson, D.P. Protruding Features of Viral Capsids are Clustered on Icosahedral Great Circles. PLoS ONE 2016, 11, 1–22. doi:10.1371/journal.pone.0152319.

15. Caspar, D.L.; Klug, A. Physical Principles in the Construction of Regular Viruses. Cold Spring Harbor Symposia on Quantitative Biology 1962, 27, 1–24. doi:10.1101/sqb.1962.027.001.005.

16. Waltmann, C.; Asor, R.; Raviv, U.; Olvera de la Cruz, M. Assembly and Stability of Simian Virus 40 Polymorphs. ACS nano 2020, 14, 4430–4443. doi:10.1021/acsnano.9b10004.

17. Prasad, B.V.V.; Schmid, M.F. Principles of Virus Structural Organization; 2012; pp. 17–47. doi:10.1007/978-1-4614-0980-9.

18. Louten, J. Virus Structure and Classification; Academic Press, 2016; pp. 19–29.

19. Baltimore, D. Expression of animal virus genomes. Bacteriology Reviews 1971, 35, 235–241.

20. Mannige, R.V.; Brooks III, C.L. Periodic table of virus capsids: Implications for natural selection and design. PLoS ONE 2010, 5, 1–7. doi:10.1371/journal.pone.0009423.

21. Wynne, S.A.; Crowther, R.A.; Leslie, A.G.W. The Crystal Structure of the Human Hepatitis B Virus Capsid. Molecular Cell 1999, 3, 771–780. doi:10.1016/S1097-2765(01)80009-5.

22. Hadden, J.A.; Perilla, J.R.; Schlicksup, C.J.; Venkatakrishnan, B.; Zlotnick, A.; Schulten, K. All-atom molecular dynamics of the HBV capsid reveals insights into biological function and cryo-EM resolution limits. eLife 2018, 7, 1–27. doi:10.7554/eLife.32478.

23. Hartman, E.C.; Jakobson, C.M.; Favor, A.H.; Lobba, M.J.; Álvarez-Benedicto, E.; Francis, M.B.; Tullman-Ercek, D. Quantitative Characterization of All Single Amino Acid Variants of a Viral Capsid-Based Drug Delivery Vehicle. Nature Communications 2018, 9, 1–11. doi:10.1038/s41467-018-03783-y.

24. Hartman, E.C.; Lobba, M.J.; Favor, A.H.; Robinson, S.A.; Francis, M.B.; Tullman-Ercek, D. Experimental Evaluation of Coevolution in a Self-Assembling Particle. Biochemistry 2019, 58, 1527–1538. doi:10.1021/acs.biochem.8b00948.

25. Venkatakrishnan, B.; Zlotnick, A. The Structural Biology of Hepatitis B Virus: Form and Function. Annual Review of Virology 2016, 3, 429–451. doi:10.1146/annurev-virology-110615-042238.

26. Nair, S.; Zlotnick, A. Asymmetric Modification of Hepatitis B Virus (HBV) Genomes by an Endogenous Cytidine Deaminase inside HBV Cores Informs a Model of Reverse Transcription. Journal of Virology 2018, 92, 1–15. doi:10.1128/jvi.02190-17.

27. Greber, B.J.; Sutter, M.; Kerfeld, C.A. The plasticity of molecular interactions governs bacterial microcompartment shell assembly. Structure 2019, 27, 749–763. doi:10.1016/j.str.2019.01.017.The.

28. El Omari, K.; Sutton, G.; Ravantti, J.J.; Zhang, H.; Walter, T.S.; Grimes, J.M.; Bamford, D.H.; Stuart, D.I.; Mancini, E.J. Plate tectonics of virus shell assembly and reorganization in phage φ8, a distant relative of mammalian reoviruses. Structure 2013, 21, 1384–1395. doi:10.1016/j.str.2013.06.017.

29. Hadfield, A.T.; Lee, W.M.; Zhao, R.; Oliveira, M.A.; Minor, I.; Rueckert, R.R.; Rossmann, M.G. The refined structure of human rhinovirus 16 at 2.15 Å resolution: Implications for the viral life cycle. Structure 1997, 5, 427–441. doi:10.1016/S0969-2126(97)00199-8.

30. McKenna, R.; Bowman, B.R.; Ilag, L.L.; Rossmann, M.G.; Fane, B.A. Atomic structure of the degraded procapsid particle of the bacteriophage G4: Induced structural changes in the presence of calcium ions and functional implications. Journal of Molecular Biology 1996, 256, 736–750. doi:10.1006/jmbi.1996.0121.

31. Golmohammadi, R.; Valegard, K.; Fridborg, K.; Liljas, L. The Refined Structure of Bacteriophage MS2 at 2·8 Å Resolution. Journal of Molecular Biology 1993, 234, 620–639. doi:10.1006/jmbi.1993.1616.

32. Erickson, J.W.; Silva, A.M.; Murthy, M.R.; Fita, I.; Rossmann, M.G. The structure of a T = 1 icosahedral empty particle from southern bean mosaic virus. Science 1985, 229, 625–629. doi:10.1126/science.4023701.

33. Sangita, V.; Satheshkumar, P.S.; Savithri, H.S.; Murthy, M.R.N. Structure of a mutant \ it T = 1 capsid of Sesbania mosaic virus: role of water molecules in capsid architecture and integrity. Acta Crystallographica Section D 2005, 61, 1406–1412.

34. Acton, O.; Grant, T.; Nicastro, G.; Ball, N.J.; Goldstone, D.C.; Robertson, L.E.; Sader, K.; Nans, A.; Ramos, A.; Stoye, J.P.; Taylor, I.A.; Rosenthal, P.B. Structural basis for Fullerene geometry in a human endogenous retrovirus capsid. Nature Communications 2019, 10, 1–13. doi:10.1038/s41467-019-13786-y.

35. Ilyas, M.; Mietzsch, M.; Kailasan, S.; Väisänen, E.; Luo, M.; Chipman, P.; Smith, J.K.; Kurian, J.; Sousa, D.; McKenna, R.; Söderlund-Venermo, M.; Agbandje-Mckenna, M. Atomic resolution structures of human bufaviruses determined by cryo-electron microscopy. Viruses 2018, 10. doi:10.3390/v10010022.

36. Guan, J.; Bywaters, S.M.; Brendle, S.A.; Ashley, R.E.; Makhov, A.M.; Conway, J.F.; Christensen, N.D.; Hafenstein, S. Cryoelectron Microscopy Maps of Human Papillomavirus 16 Reveal L2 Densities and Heparin Binding Site. Structure 2017, 25, 253–263. doi:10.1016/j.str.2016.12.001.

37. Wolf, M.; Garcea, R.L.; Grigorieff, N.; Harrison, S.C. Subunit interactions in bovine papillomavirus. Proceedings of the National Academy of Sciences of the United States of America 2010, 107, 6298–6303. doi:10.1073/pnas.0914604107.

38. Stehle, T.; Gamblin, S.J.; Yan, Y.; Harrison, S.C. The structure of simian virus 40 refined at 3.1 Å resolution. Structure 1996, 4, 165–182. doi:10.1016/S0969-2126(96)00020-2.

39. Lindner, J.M.; Cornacchione, V.; Sathe, A.; Be, C.; Srinivas, H.; Riquet, E.; Leber, X.C.; Hein, A.; Wrobel, M.B.; Scharenberg, M.; Pietzonka, T.; Wiesmann, C.; Abend, J.; Traggiai, E. Human Memory B Cells Harbor Diverse Cross-Neutralizing Antibodies against BK and JC Polyomaviruses. Immunity 2019, 50, 668–676.e5. doi:10.1016/j.immuni.2019.02.003.

40. Yadav, R.; Zhai, L.; Tumban, E. Virus-like Particle-Based L2 Vaccines against HPVs: Where Are We Today? Viruses 2020, 12, 1–17. doi:10.7765/9780719098451.00007.

41. Mietzsch, M.; Pénzes, J.J.; Agbandje-Mckenna, M. Twenty-five years of structural parvovirology. Viruses 2019, 11. doi:10.3390/v11040362.

42. Callaway, H.M.; Subramanian, S.; Urbina, C.A.; Barnard, K.N.; Dick, R.A.; Bator, C.M.; Hafenstein, S.L.; Gifford, R.J.; Parrish, C.R. Examination and Reconstruction of Three Ancient Endogenous Parvovirus Capsid Protein Gene Remnants Found in Rodent Genomes. Journal of Virology 2019, 93, 1–14. doi:10.1128/jvi.01542-18.

43. Halder, S.; Nam, H.J.; Govindasamy, L.; Vogel, M.; Dinsart, C.; Salome, N.; McKenna, R.; Agbandje-McKenna, M. Structural Characterization of H-1 Parvovirus: Comparison of Infectious Virions to Empty Capsids. Journal of Virology 2013, 87, 5128–5140. doi:10.1128/jvi.03416-12.

44. Tan, Y.Z.; Aiyer, S.; Mietzsch, M.; Hull, J.A.; McKenna, R.; Grieger, J.; Samulski, R.J.; Baker, T.S.; Agbandje-McKenna, M.; Lyumkis, D. Sub-2 Å Ewald curvature corrected structure of an AAV2 capsid variant. Nature Communications 2018, 9, 1–11. doi:10.1038/s41467-018-06076-6.

45. Meyer, N.L.; Hu, G.; Davulcu, O.; Xie, Q.; Noble, A.J.; Yoshioka, C.; Gingerich, D.S.; Trzynka, A.; David, L.; Stagg, S.M.; Chapman, M.S. Structure of the gene therapy vector, adeno-associated virus with its cell receptor, aavr. eLife 2019, 8, 1–24. doi:10.7554/eLife.44707.

46. Burmeister, W.P.; Buisson, M.; Estrozi, L.F.; Schoehn, G.; Billet, O.; Hannas, Z.; Sigoillot, C.; Poulet, H. Structure determination of feline calicivirus virus-like particles in the context of a pseudo-octahedral arrangement. PLoS ONE 2015, 10, 1–15. doi:10.1371/journal.pone.0119289.

47. Duquerroy, S.; Da Costa, B.; Henry, C.; Vigouroux, A.; Libersou, S.; Lepault, J.; Navaza, J.; Delmas, B.; Rey, F.A. The picobirnavirus crystal structure provides functional insights into virion assembly and cell entry. EMBO Journal 2009, 28, 1655–1665. doi:10.1038/emboj.2009.109.

48. Wada, Y.; Tanaka, H.; Yamashita, E.; Kubo, C.; Ichiki-Uehara, T.; Nakazono-Nagaoka, E.; Omura, T.; Tsukihara, T. The structure of melon necrotic spot virus determined at 2.8 Å resolution. Acta Crystallographica Section F: Structural Biology and Crystallization Communications 2008, 64, 8–13. doi:10.1107/S1744309107066481.

49. Guu, T.S.Y.; Liu, Z.; Ye, Q.; Mata, D.A.; Li, K.; Yin, C.; Zhang, J.; Tao, Y.J. Structure of the hepatitis E virus-like particle suggests mechanisms for virus assembly and receptor binding. Proceedings of the National Academy of Sciences 2009, 106, 12992–12997. doi:10.1073/pnas.0904848106.

50. Chen, N.C.; Yoshimura, M.; Guan, H.H.; Wang, T.Y.; Misumi, Y.; Lin, C.C.; Chuankhayan, P.; Nakagawa, A.; Chan, S.I.; Tsukihara, T.; Chen, T.Y.; Chen, C.J. Crystal Structures of a Piscine Betanodavirus: Mechanisms of Capsid Assembly and Viral Infection. PLoS Pathogens 2015, 11, 1–25. doi:10.1371/journal.ppat.1005203.

51. Pan, J.; Dong, L.; Lin, L.; Ochoa, W.F.; Sinkovits, R.S.; Havens, W.M.; Nibert, M.L.; Baker, T.S.; Ghabrial, S.A.; Tao, Y.J. Atomic structure reveals the unique capsid organization of a dsRNA virus. Proceedings of the National Academy of Sciences of the United States of America 2009, 106, 4225–4230. doi:10.1073/pnas.0812071106.

52. Jung, J.; Grant, T.; Thomas, D.R.; Diehnelt, C.W.; Grigorieff, N.; Joshua-Tor, L. High-resolution cryo-EM structures of outbreak strain human norovirus shells reveal size variations. Proceedings of the National Academy of Sciences of the United States of America 2019, 116, 12828–12832. doi:10.1073/pnas.1903562116.

53. Madhumati, S.; Long, F.; Miller, A.; Klose, T.; Buda, G.; Sun, L.; Kuhn, R.; Rossmann, M.G. Refinement and Analysis of the Mature Zika Virus Cryo-EM Structure at 3.1 Å Resolution. Physiology & behavior 2018, 26, 1169–1177. doi:10.1016/j.str.2018.05.006.Refinement.

54. Wolf, Y.I.; Kazlauskas, D.; Iranzo, J.; Lucía-Sanz, A.; Kuhn, J.H.; Krupovic, M.; Dolja, V.V.; Koonin, E.V. Origins and evolution of the global RNA virome. mBio 2018, 9. doi:10.1128/mBio.02329-18.

